# Synergistic activity of Nanog, Pou5f3, and Sox19b establishes chromatin accessibility and developmental competence in a context-dependent manner

**DOI:** 10.1101/2020.09.01.278796

**Authors:** Liyun Miao, Yin Tang, Ashley R. Bonneau, Shun Hang Chan, Mina L. Kojima, Mark E. Pownall, Charles E. Vejnar, Antonio J. Giraldez

## Abstract

Genome-wide chromatin reprogramming is a fundamental requirement for establishing developmental competence in the newly-formed zygote. In zebrafish, Nanog, Pou5f3 and Sox19b play partially redundant roles in zygotic genome activation, however their interplay in establishing chromatin competency, the context in which they do so and their mechanism of action remain poorly defined. Here, we generated a triple maternal-zygotic *nanog*^-/-^;*pou5f3*^-/-^;*sox19b*^-/-^ mutant and assessed the causal relationship between transcription factor (TF) occupancy, chromatin accessibility and genome activation. Analyses of this triple mutant and combinatorial rescues revealed highly synergistic and context-dependent activity of Nanog, Pou5f3, and Sox19b (NPS) in establishing chromatin competency at >50% of active enhancers. Motif analysis revealed a network of TFs that depend on NPS for establishing chromatin accessibility, including the endodermal determinant Eomesa, whose binding we show is regulated by NPS pioneer-like activity. Finally, we demonstrated that NPS play an essential role in establishing H3K27ac and H3K18ac at enhancers and promoters, and that their function in transcriptional activation can be bypassed by targeted recruitment of histone acetyltransferases to individual genes. Altogether, our findings reveal a large network of TFs that function to establish developmental competency, many of which depend on the synergistic and highly context-dependent role of NPS in establishing chromatin accessibility and regulating histone acetylation in order to activate the genome.

---

**T**he newly formed embryo is transcriptionally silent and undergoes genome-wide chromatin reprogramming to initiate zygotic genome activation (ZGA) and gain transient totipotency (Iwafuchi-Doi and Zaret, 2014; Ladstätter and Tachibana, 2019). This critical step in development marks the transfer of regulatory control from the maternal program to the zygotic program and is known as the maternal-to-zygotic transition (MZT) (Giraldez, 2010; Tadros and Lipshitz, 2009; Vastenhouw et al., 2019; Yartseva and Giraldez, 2015). Individual transcription factors (TFs) regulating this process have been identified, for example Zelda and GAGA Factor (GAF) in *Drosophila (Gaskill et al., 2020; Harrison et al., 2011; Liang et al., 2008; Nien et al., 2011; Schulz et al., 2015; Sun et al., 2015); Nanog, Pou5f3 (OCT4 homolog), and SoxB1 family in zebrafish* (Joseph et al., 2017; Lee et al., 2013; Leich-senring et al., 2013); and Dppa2, Dppa4, and NFYA in mice (Eckersley-Maslin et al., 2019; Lu et al., 2016). OCT4 contributes to ZGA in human embryos but not in mice (Foygel et al., 2008; Gao et al., 2018; Le Bin et al., 2014; Wu et al., 2013). DUX4 proteins, on the other hand, have been shown to regulate ZGA in both mice and humans (De Iaco et al., 2017; Hendrickson et al., 2017; Whiddon et al., 2017). While these studies have identified a small subset of TFs that function during ZGA, the larger regulatory network of TFs that orchestrate chromatin remodeling, their interdependency and how they trigger genome activation are still poorly understood.

The local chromatin landscape undergoes substantial reorganization during the MZT (Eckersley-Maslin et al., 2018). An increase in chromatin accessibility coincides with genome activation and these newly accessible regions are associated with zygotic transcription (Blythe and Wieschaus, 2016; Gao et al., 2018; Li et al., 2018; Liu et al., 2018; Lu et al., 2016; Vastenhouw et al., 2019; Wu et al., 2016; Wu et al., 2018). However, the regulatory mechanisms underlying chromatin remodeling to gain transcriptional competency during the MZT remain largely unclear. Pioneer factors can bind condensed, nucleosomal DNA to promote chromatin accessibility, thereby allowing other TFs to bind and creating a permissive state conducive to gene expression (Iwafuchi-Doi, 2019; Iwafuchi-Doi and Zaret, 2016). Recent studies have shown that ZGA activators, such as Zelda in *Drosophila*, FoxH1 in *Xenopus*, OCT4 in human, and NFYA in mice, exhibit pioneering activity (Cirillo et al., 2002; Iwafuchi-Doi and Zaret, 2014) as they initiate chromatin opening and mediate the binding of other TFs to chromatin (Charney et al., 2017; Gao et al., 2018; Lu et al., 2016; Schulz et al., 2015; Sun et al., 2015). However, the extent of their binding is much broader than the extent to which they regulate chromatin accessibility; hence, the genomic context that determines pioneer function *in vivo* is largely unclear. Additionally, many enhancers are co-occupied by multiple TFs (Chen et al., 2008; Struhl, 2001), indicating functional redundancy. For example, Nanog, Pou5f3, and SoxB1 family co-bind many regions in the genome, and play combinatorial roles in the regulation of ZGA in zebrafish (Lee et al., 2013; Leichsenring et al., 2013). Loss of a single one of these factors leads to decreased chromatin accessibility at only a subset of their bound regions, suggesting that their pioneering activity is context-dependent and/or may be regulated by additional factors in vivo (Charney et al., 2017; Gao et al., 2020; Pálfy et al., 2020; Veil et al., 2019b). What determines this context has not yet been defined, and whether they serve as pioneer factors for other TFs to establish chromatin competency across the genome is unknown.

Reprogramming of the genome during the MZT coincides with major epigenetic remodeling of the chromatin including remodeling of the DNA methylation, incorporation of specific histone variants, and histone modifications (Eck-ersley-Maslin et al., 2018; Murphy et al., 2018; Vastenhouw et al., 2019). Histone H3 lysine 4 trimethylation (H3K4me3) often marks active promoters, while Histone H3 lysine 27 trimethylation (H3K27me3) is correlated with inactive loci; and the presence of both marks increase dramatically during the MZT (Vastenhouw et al., 2019; Vastenhouw et al., 2010; Xia et al., 2019; Zhang et al., 2018; Zhu et al., 2019). Histone H3 lysine 27 acetylation (H3K27ac) correlates with regions of accessible chromatin and often marks active enhancers and promoters (Creyghton et al., 2010; Wang et al., 2008). H3K27ac precedes ZGA in zebrafish (Chan et al., 2019; Sato et al., 2019; Zhang et al., 2018). In addition to H3K27ac, acetylation of histone H3 lysine 18 (H3K18ac) and histone H4 lysine 8 (H4K8ac) are enriched at zygotic genes prior to ZGA in *Drosophila* (Li et al., 2014), and overexpression of p300 (a histone acetyltransferase) and BRD4 (a reader of histone acetyl lysine) in zebrafish prematurely activates zygotic transcription, suggesting a potential regulatory role for histone acetylation in ZGA (Chan et al., 2019). Additionally, H4K16ac is present in oocytes and has been shown to regulate local chromatin accessibility before ZGA in *Drosophila* (Samata et al., 2020). Acetylated chromatin is often relatively more accessible (Hebbes et al., 1994; Krajewski and Becker, 1998; Lee et al., 1993; Simpson, 1978) and biochemical studies have revealed that acetylation reduces the affinity between DNA and nucleosomes (Cary et al., 1982; Garcia-Ramirez et al., 1995; Tse et al., 1998; Yau et al., 1982). However, the causal relationship between histone acetylation and chromatin accessibility *in vivo* remains unclear.

Here, we performed combinatorial genetic knockout and rescue analyses using a triple maternal-zygotic *nanog*^**-/-**^; *pou5f3*^**-/-**^; *sox19b*^**-/-**^ mutant and assess the causal relationship between TF occupancy, chromatin accessibility, and transcriptional activation during the MZT. Our study revealed that Nanog, Pou5f3, and Sox19b—the maternally loaded member of the soxB1 family (Lee et al., 2013; Pálfy et al., 2020)— interact synergistically to achieve developmental reprogramming by opening >50% of active enhancers. Motif analysis revealed a set of TFs that overlap accessible regions during genome activation and their dependency on NPS pioneer activity. We further identified sequence and nucleosome features that facilitates pioneer activity *in vivo*. Finally, we demonstrated that NPS play an essential role in establishing H3K27ac at enhancers and promoters, and their function in transcriptional activation can be bypassed by targeted recruitment of histone acetyltransferases to individual genes. Altogether, our findings reveal a critical regulatory role for the genomic context in the regulation of Nanog, Pou5f3, and Sox19b binding during the MZT, which in turn establishes transcriptional competence and facilitates subsequent binding of developmentally important TFs. We further show how NPS regulate enhancer accessibility and trigger genome activation through histone acetylation and recruitment of RNA polymerase II, delineating a sequence of events in genome activation.

## Results

### Nanog, Pou5f3, and Sox19b regulate chromatin accessibility at many genomic regions, particularly at active enhancers

Genome-wide chromatin reprogramming during the MZT generates accessible regions in the chromatin that are important for regulating subsequent developmental programs. To gain a global view of chromatin accessibility during this transition, we performed Omni-ATAC (Buenrostro et al., 2013; Corces et al., 2017) across three biological replicates during the MZT, at 4 hours post-fertilization (hpf). We identified 55,970 high-confidence accessible regions (Figures S1A and S1B) including 17,993 at promoters (defined as the region within 500 bp of the transcription start site (TSS) of annotated transcripts) and a further 16,142 accessible regions marked by H3K27ac, suggestive of active enhancers (Figure 1A). We next sought to determine what fraction of the accessible chromatin is bound by Nanog, Pou5f3, and Sox19b. To this end, we analyzed newly generated ChIP-seq data for Pou5f3 and Sox19b, as well as previously published Nanog ChIP-seq data (Xu et al., 2012). Among all accessible regions, 26,441 (47%) were occupied by at least one of these factors. 12,694 regions (23%) were co-occupied by two or more of these factors (Figure 1B). 10,636 of these regions overlapped with H3K27ac, suggestive of active enhancers or promoters (Figure 1C). These data reveal that Nanog, Pou5f3, and Sox19b, either alone or in combination, occupy a significant fraction of the accessible chromatin, particularly at active enhancers or promoters.

**Figure 1.**
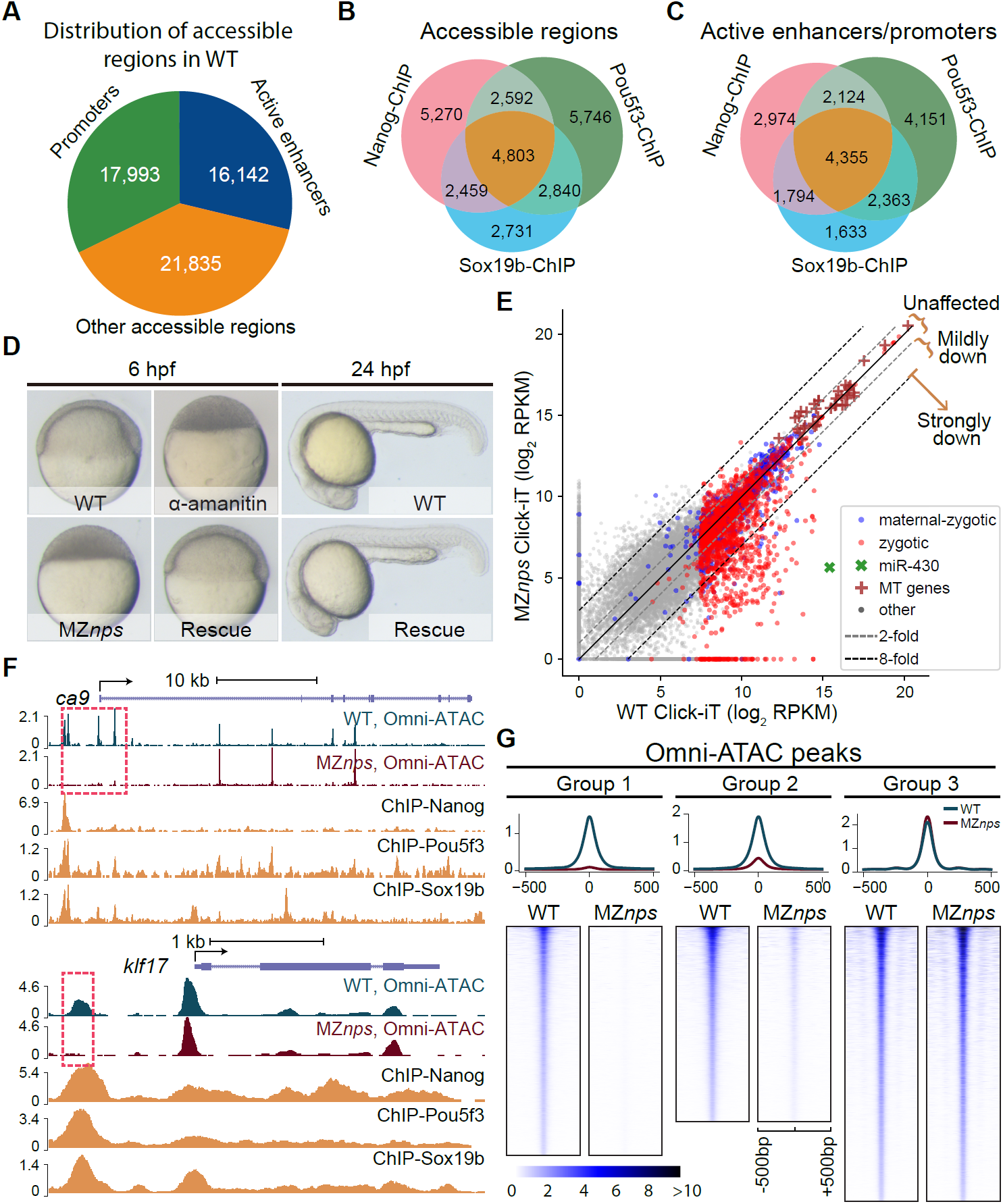
Nanog, Pou5f3, and Sox19b (NPS) regulates chromatin accessibility at a significant fraction of open regions. (A) Distribution of accessible regions among putative promoters, active enhancers, and other accessible regions in wild-type (WT) embryos at 4 hpf. (B) Venn diagram showing significant overlap of NPS binding, as assayed by ChIP-seq, at all accessible regions (*P* < 1×10^-100^ Fisher’s exact test) (C) Venn diagram showing a significant overlap between ChIP-seq peaks for NPS within accessible regions positive for H3K27ac (putative active promoters and enhancers) (*P* < 1×10^−100^ Fisher’s exact test). (D) Maternal and Zygotic mutant for *nanog;pou5f3;sox19b* (MZ*nps*) arrested before gastrulation. At 6 hpf, WT embryos had developed to the shield stage, while MZ*nps* embryos resemble embryos treated with α-amanitin and arrest at sphere stage. Co-injection of *nanog* and *pou5f3* mRNAs (Rescue) rescued development of the MZ*nps* embryos at 6 hpf and 24 hpf. (E) Biplots comparing the nascent transcriptome of MZ*nps* embryos with that of WT embryos using click-it seq. MT genes: mitochondrial genes. See methods for the definition of zygotic and maternally zygotic genes. *Strongly down*: Zygotic genes with nascent transcription that are downregulated ≥ 8-fold in MZ*nps* compared to WT embryos. *Mildly down*: Zygotic genes with nascent transcription that are downregulated between 8- and 2-fold in MZ*nps* compared to WT embryos. *Unaffected*: Zygotic genes with transcription that is not substantially changed (< 2-fold) in MZ*nps* compared to WT embryos. (F) Representative genomic tracks of Omni-ATAC in WT and MZ*nps* embryos, and ChIP-seq for the indicated factors in WT embryos. Signal intensity in RPM (reads per million). The dashed box highlights accessible regions in wild-type (WT) that are inaccessible to ATAC in the MZ*nps* embryos. (G) Line plots (above) and heatmaps (below) of Omni-ATAC signal at accessible regions in WT and MZ*nps* embryos. Group 1 include 7,813 regions without detectable accessibility in MZ*nps* embryos (see Methods). Group 2 include 6,665 regions with decreased accessibility in MZ*nps* embryos. Group 3 include 9,371 regions with similar accessibility WT and MZ*nps* embryos. Heatmaps are centered at the summit of Omni-ATAC peaks with 500 bp on both sides and are ranked from high to low according to the average intensity of the Omni - ATAC signal in WT.

To determine the causal relationship between Nanog, Pou5f3, and Sox19b and the establishment of chromatin accessibility, we generated a triple maternal-zygotic (MZ) mutant for *nanog*^**-/-**^;*pou5f3*^**-/-**^;*sox19b*^**-/-**^ (MZ*nps*; hereafter we use “NPS” to refer to Nanog, Pou5f3, and Sox19b) (Figure S1C). MZ mutants are both deficient in zygotic gene expression and also lack any maternally-inherited protein. All MZ*nps* embryos did not gastrulate and were arrested at the sphere stage (4 hpf) (Figure 1D), a phenotype that is reminiscent of a block in zygotic development (Kane et al., 1996). Analysis of the MZ*nps* nascent transcriptome using Click-iT-seq (Chan et al., 2019) revealed that 37% of zygotic genes (822 out of 2,240) (Figures 1E and S1D) were downregulated more than twofold compared to wild-type embryos. Consistent with this, RNA polymerase II (Pol II) ChIP-seq revealed a concomitant loss of Pol II occupancy across the gene body and promoters of these genes (Figure S1E), suggesting a critical role for NPS in the recruitment of Pol II, rather than in the release of paused Pol II. Gene ontology analysis (Ge et al., 2019; Mi et al., 2019a; Mi et al., 2019b) revealed that NPS targets are enriched in gene-specific transcriptional regulator and in developmental processes, including embryonic morphogenesis, formation of primary germ layers, and cell fate commitment (Figures S1F and S1G).

Next, we identified 14,478 regions (out of 55,970) that had a significant decrease in accessibility in MZ*nps* compared to wild-type embryos using DESeq2 (Love et al., 2014) (FDR < 0.01; Figures 1F, 1G and S1H). Among these, 7,813 lost accessibility (Group 1) and were completely dependent on NPS, while 6,665 regions (Group 2) remained partially accessible in MZ*nps* embryos, suggesting that other factors co-bind with NPS and function to displace nucleosomes at these regions. Over half of active enhancers (8,274 out of 16,142) showed decreased chromatin accessibility, which correlated with the loss of transcription for zygotic genes (Pearson correlation, Student’s ttest; *r* = 0.49, *P* = 8.8 × 10^−60^) (Figures 2A, S2A and S2B). In contrast, only 5.1% (914 out of 17,993) of promoters showed a significant decrease in accessibility in MZ*nps* embryos (Figure 2A). Promoter accessibility was maintained even in genes that were downregulated in MZ*nps* embryos despite the loss of Pol II ChIP-seq signal (Figures 2B and S2C). Altogether, these results indicate that NPS are responsible for establishing chromatin accessibility at the majority of active enhancers during the MZT, while promoter accessibility is largely regulated by other factors independently of NPS and Pol II. Furthermore, these data provide an entry point to understand what factors co-regulate chromatin accessibility and ZGA, both together with and independent of NPS, and the causal relationship be tween their binding to chromatin and the epigenetic remodeling that regulates genome activation.

**Figure 2.**
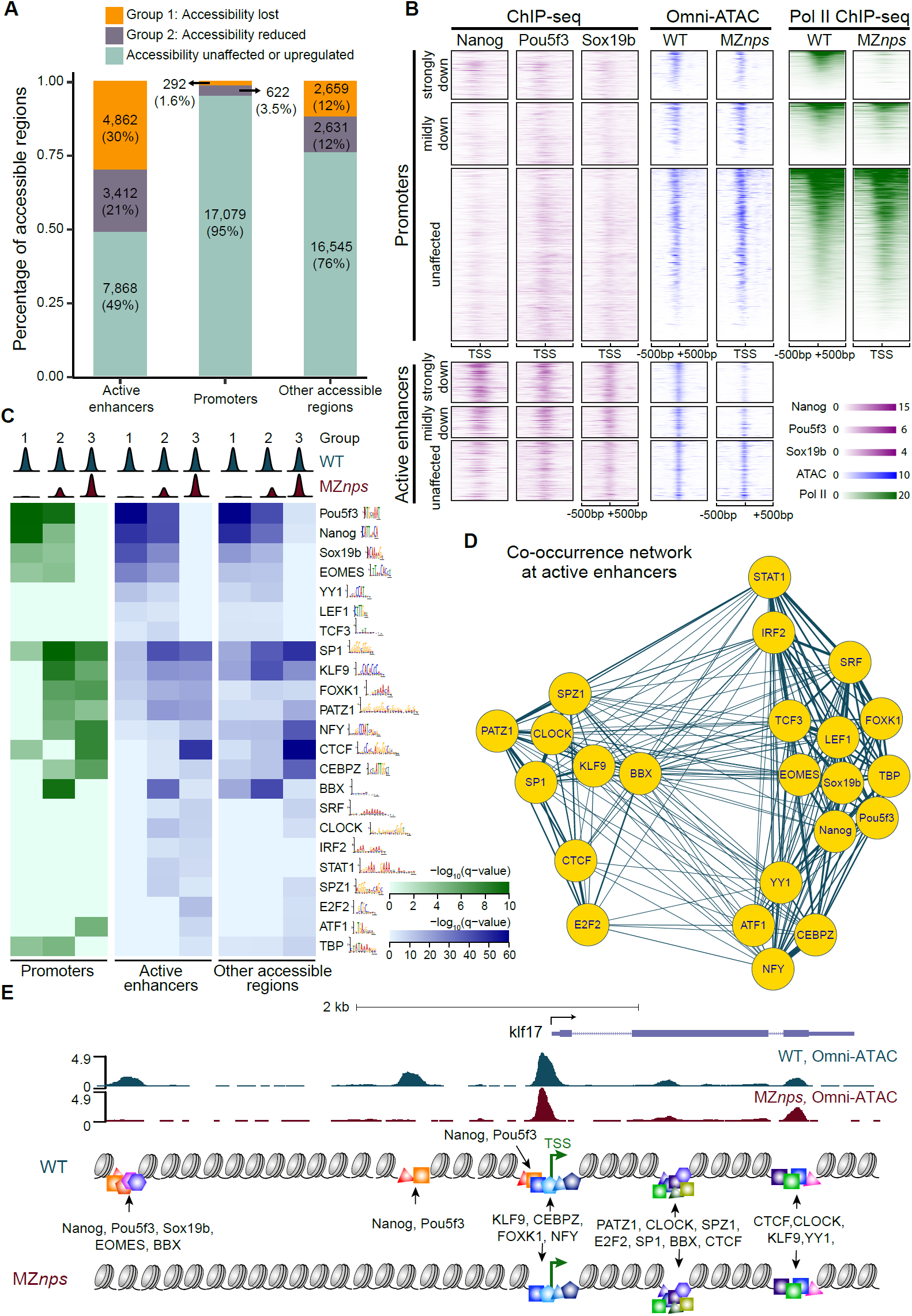
Identification of TF motifs overlapping chromatin accessibility and their dependence on Nanog, Pou5f3, and Sox19b. (A) Stacked bar plot showing the percentage of accessible regions dependent on NPS (Group 1, 2), and accessible regions that are unaffected or upregulated in MZ*nps* across different genomic locations. Active enhancers contained the highest percentage of regions with reduced accessibility in MZ*nps*, and promoters were mostly unaffected. (B) Heatmaps showing accessibility (Omni - ATAC), Pol II binding and NPS ChIP-seq binding intensities at promoter and active enhancer regions for different categories of zygotic genes in wild-type (WT) and MZ*nps* embryos. Heatmaps are centered at the TSS (promoters) and the summit of the Omni-ATAC peak (active enhancers) with 500 bp on both sides and ranked according to the average intensity of Pol II binding within the promoter region of the zygotic genes in WT embryos from high to low. Strongly down, mildly down and unaffected refers to the effect on zygotic transcription in MZ*nps* as described in Figure 1E. For strongly downregulated genes, Pol II levels near the TSS are depleted in MZ*nps* but accessibility remains. In contrast, active enhancers for strongly downregulated genes show the higher binding intensity for NPS and the strongest reduction in accessibility. (C) Enrichment of known sequence motifs of TFs at enhancers and promoters within the three groups of accessible regions from Figure 1G. (D) Co-occurrence of the enriched sequence motifs of TFs at active enhancers in (C). The closer distance and thicker connection between nodes represent higher co-occurrence between the sequence motifs of the TFs in the network. (E) Diagram illustrating the sequence motifs of TFs located at the accessible regions near the *klf17* gene. The locations of the sequence motifs agree with the enrichment of these motifs at different locations in (C).

### Identification of the TF network regulating chromatin accessibility during the MZT and their dependency of NPS

The critical nature of Nanog, Pou5f3, and Sox19b in establishing chromatin accessibility during the MZT is reminiscent of the pioneering activity of some of their mammalian homologs (Heurtier et al., 2019; Zaret and Carroll, 2011). We reasoned that NPS may function in a similar manner to facilitate the binding of a network of TFs necessary for genome activation during the MZT. To identify the TFs regulating genome activation and their dependency on NPS, we first determined those TFs that are maternally expressed and translated pre-ZGA using RNA-seq and ribosome footprinting data at 2 hpf (Bazzini et al., 2012). Then, based on the loss of accessibility in MZ*nps* embryos, we sub-divided enhancers, promoters, and the other accessible regions into three groups and analyzed motif enrichment for different TFs using MEME suite (McLeay and Bailey, 2010). Because Group 1 completely lost chromatin accessibility in MZ*nps* embryos, we hypothesized that NPS are the key regulators of Group 1 regions and possibly function in a pioneer-like role to open these regions and establish a permissive environment for other TFs to bind. Group 2 showed reduced accessibility in MZ*nps* embryos, therefore we hypothesized that NPS are partially responsible for establishing accessibility at these regions, but other factors are able to initiate chromatin opening independent of NPS. Group 3 accessibility was not significantly affected in MZ*nps* embryos, and therefore we hypothesized that other factors must regulate accessibility in these regions.

Consistent with a dependency on NPS, motif enrichment analysis revealed that Groups 1 and 2 were enriched for Nanog, Pou5f3, and Sox19b motifs (Figure 2C). Interestingly, we also observed a significant enrichment for other TF motifs, including EOMES, YY1, LEF1, TCF3, and TBP. These factors were enriched in both groups, suggesting that in some regions NPS is required to establish chromatin accessibility (Group1), while in other regions they are able to establish accessibility independent of NPS (Group 2). Factors such as BBX, PATZ1, and KLF9 were more enriched in Group 2, suggesting that these factors co-occupy Group 2 regions with NPS but can establish chromatin opening independent of NPS. Other factors were significantly enriched in regions that establish accessibility independent of NPS, (Group 2 and 3), including SRF, CLOCK, IRF2, STAT1, SPZ1, E2F2, and ATF1. We observed that accessible promoters independent of NPS (Group 3) were significantly enriched for SP1, KLF9, FOXK1, PATZ1, NFY, CEBPZ, andATF1 motifs (Figure 2C). Interestingly, the factors that were dependent on NPS were significantly enriched for developmental functions (5/7; Hypergeometric test, *P* = 3.9 × 10^−5^) while most (12/13) TFs that were independent of NPS were related to housekeeping functions (Eisenberg and Levanon, 2013; Murphy et al., 2018), suggesting that the NPS-associated regulatory network primarily orchestrates the developmental program.

Combinatorial TF binding and/or TF redundancy are common genetic mechanisms that enable appropriate activation and function of gene regulatory networks during development (Boyer et al., 2005; Chen et al., 2008; Loh et al., 2006; Struhl, 2001; Wang et al., 2012). To identify TFs with shared regulatory elements in the genome, we analyzed the pairwise enrichment of different motifs and constructed a co-occurrence network (Figures 2D, 2E, and S2D). We observed significant co-occurrence between Nanog, Pou5f3, and Sox19b motifs at promoters and active enhancers. In addition, motifs for Eomesa, LEF1, and TBP were more often associated with NPS than with other TF motifs. In contrast, SP1, KLF9, PATZ1, SPZ1, and CLOCK were more often found together than with other TFs. Altogether, these results reveal a TF network dependent and independent on the pioneer-like function of NPS with the capacity to regulate chromatin accessibility during the MZT.

### Nanog, Pou5f3, and Sox19b establish a permissive chromatin state for Eomesa binding

Our analysis revealed several TF motifs that were enriched in regions that lost accessibility in MZ*nps* embryos, suggesting that NPS may be required to regulate their binding to the chromatin. Among these TFs, Eomesodermin (homolog of Eomesa) is a T-box transcription factor important for endoderm and mesoderm formation in vertebrates (Bjornson et al., 2005; Nelson et al., 2014). To determine if Eomesa binding to the chromatin requires the activity of NPS, we performed Eomesa ChIP-seq in wild-type and MZ*nps* embryos at 4 hpf. Although *eomesa* transcript levels are similar between wild-type and MZ*nps* embryos (Figure S3A), Eomesa binding was completely lost at 63% of Eomesa-bound regions in MZ*nps* embryos (Figures 3A). The reduction of Eomesa binding in MZ*nps* embryos correlates with reduced accessibility at those regions (Figures 3A-3C and S3B). These results suggest that the activity of NPS in opening otherwise inaccessible chromatin is crucial for the majority of Eomesa binding.

**Figure 3.**
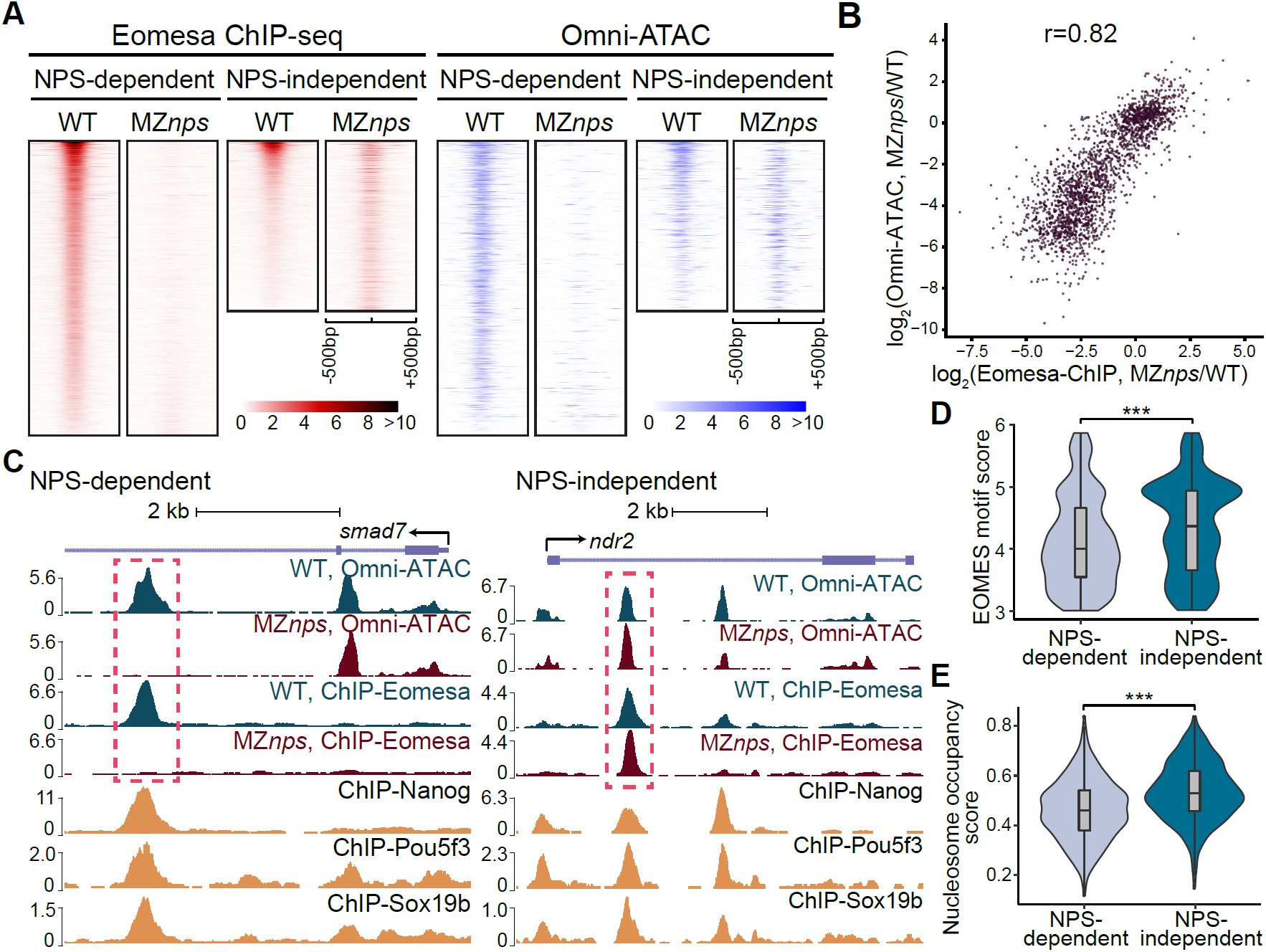
The pioneering activity of Nanog, Pou5f3, and Sox19b is required for Eomesa binding. (A) Heatmaps of Eomesa ChIP-seq and accessibility at regions where Eomesa binding was lost (NPS-dependent) or remained (NPS-independent) in MZ*nps* compared to wild-type (WT) embryos (See Methods). Heatmaps are centered on the summit of Eomesa ChIP-seq peaks with 500 bp on both sides and ranked by WT signal intensity. NPS-dependent: 1,305 regions; NPS-independent: 754 regions. (B) Biplot showing a high correlation between the reduction in Eomesa binding and accessibility in (MZ*nps*/WT) (*r* = 0.82, Pearson correlation). (C) Representative genomic tracks for NPS-dependent and NPS-independent Eomesa binding sites. NPS-dependent track: The accessible region framed within the dashed line lost both accessibility and Eomesa binding in MZ*nps* embryos. NPS-independent track: Accessibility at the region and Eomesa binding framed within the dashed line were both unaffected in MZ*nps* embryos. (D) Violin plots comparing Eomesa motif scores between NPS-dependent (n = 1117) and NPS-independent (n = 606) Eomesa binding regions. NPS-independent regions had significantly higher Eomesa motif scores than NPS dependent regions (*P* = 7.0×10^−10^, Wilcoxon rank-sum test). (E) Violin plots comparing nucleosome occupancy score between NPS-dependent (n = 1,305) and NPS-independent (n = 754) Eomesa binding regions. NPS-independent regions have significantly higher nucleosome occupancy than NPS-dependent regions (*P* = 3.7×10^−38^, Two-sample t-test). *** *P* <0.001.

Given that 37% of Eomesa-bound sites remained in the absence of Nanog, Pou5f3, and Sox19b, it is possible that Eomesa alone can initiate chromatin opening at these regions. Binding sites of pioneer factors, such as Oct4, Sox2, and Klf4 (Soufi et al., 2015), are associated with high nucleosome occupancy. Interestingly, NPS-independent Eomesa-binding regions exhibited higher EOMES motif scores (Wilcoxon rank sum test; *P* = 7.0 × 10^−10^) and higher nucleosome occupancy scores (Two-sample t-test; *P* = 3.7 × 10^−38^), which are predictive of nucleosome occupancy and positioning (Kaplan et al., 2009) (Figures 3D and 3E). While NPS-dependent Eomesa-binding regions had stronger enrichment for Nanog, Pou5f3, and Sox19b motifs, NPS-independent regions were enriched for SP1, SALL4, SMAD3, NR4A1, SRF, ZBTB18, ARID3B, E2F3, and CPEB1 motifs (Figure S3C), suggesting that one or more of these factors may be required for Eomesa binding at these regions. Altogether, these results indicate that the majority of Eomesa binding events depend on the pioneer-like activity of NPS to establish chromatin accessibility during the MZT.

### Nanog, Pou5f3, and Sox19b act both independently and cooperatively to establish chromatin accessibility

Our results indicate a significant role for NPS in establishing chromatin accessibility during the MZT. However, it is unclear which of these factors is capable of initiating chromatin opening, whether there is redundancy and/or synergy between them, and in which genomic contexts these alternatives occur. These questions are particularly relevant given the prevalence of active enhancers co-bound by all three factors during the MZT and cannot be addressed by *in vitro* experiments or single loss-of-function analyses (Cirillo et al., 2002; Pálfy et al., 2020; Veil et al., 2019a; Zaret and Carroll, 2011). To distinguish between these scenarios, we restored Nanog (N), Pou5f3 (P), or Sox19b (S) function separately and in all combinations (NP, NS, PS, NPS) in MZ*nps* embryos and assessed chromatin accessibility using Omni-ATAC in each condition at 4 hpf. To identify the physiological levels of mRNA required to restore the function of each factor, we titrated the level of mRNAs such that they rescued gastrulation and survival to adulthood without causing an over-expression phenotype. Among the 7,813 Group 1 regions where accessibility was completely lost in MZ*nps* embryos, we found that each single factor can restore chromatin accessibility at many loci (Sox19b: 201 regions; Pou5f3: 329 regions; Nanog: 2,703 regions) (Figures 4A, 4B, and S4A). While Nanog has not yet been defined as a pioneer factor *in vitro*, our results reveal that restoration of Nanog function was able to initiate chromatin opening at more regions than Pou5f3 (number of regions can be rescued by Nanog versus by Pou5f3; Fisher’s exact test, *P* < 10^−100^) and Sox19b (number of regions can be rescued by Nanog versus by Sox19b; Fisher’s exact test, *P* < 10^−100^). This is consistent with our previous observations that Nanog loss-of-function produced a stronger effect on the transcriptome than Pou5f3 or Sox19b loss-of-function (Lee et al., 2013). The activity of these proteins in establishing chromatin accessibility was concentration dependent, as we found that a higher concentration of Sox19b increased the number of regions with rescued accessibility (Figures 4A, 4B, and S4A). Although the single factor rescue was effective in restoring accessibility across many loci, co-injection of two or three factors simultaneously rescued more regions than any single factor alone (Figure 4B), suggesting an additive effect between the factors. Altogether, these results indicate that Nanog, Pou5f3, or Sox19b have pioneer-like functions *in vivo* to initiate chromatin opening.

**Figure 4.**
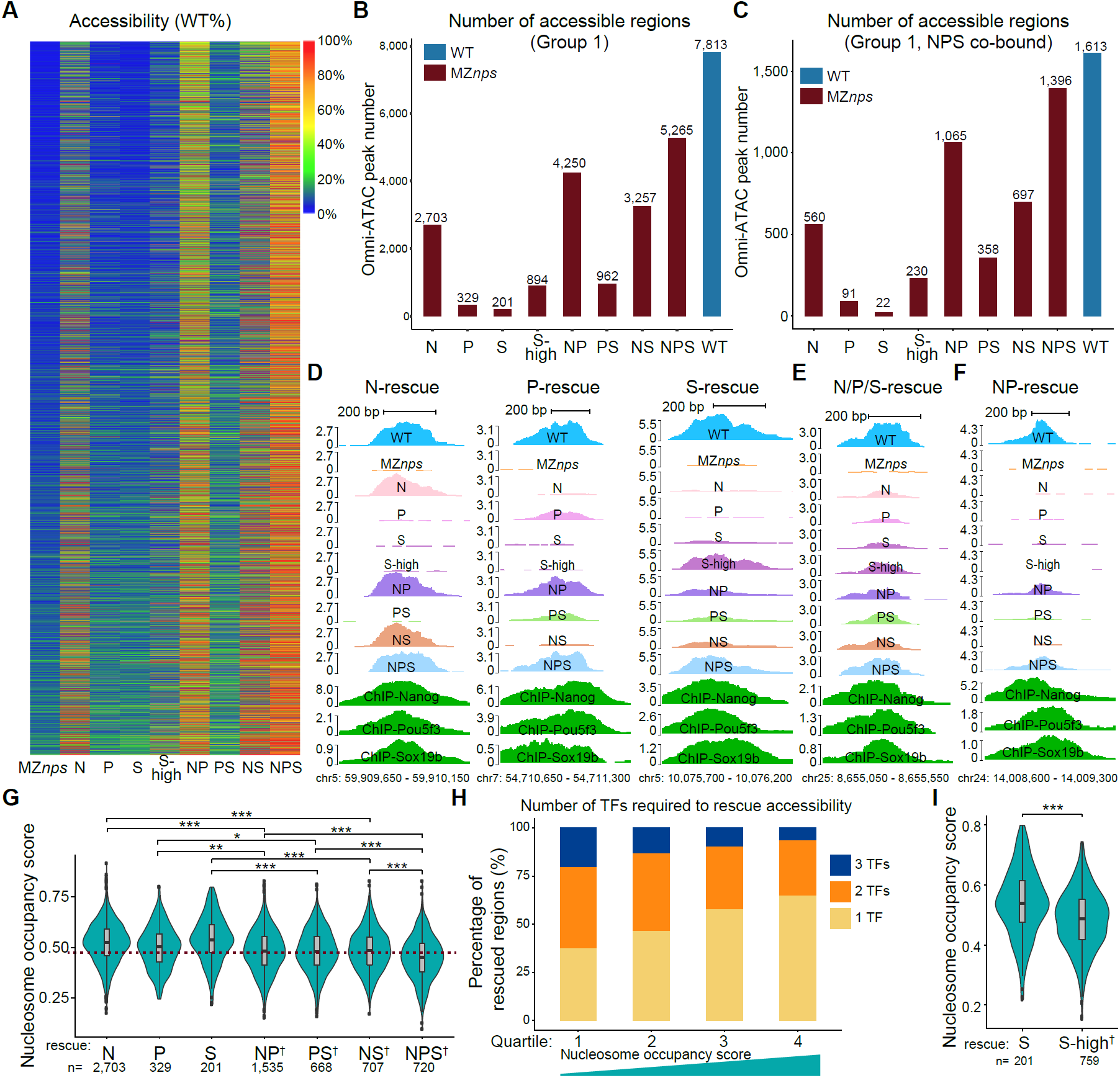
Nanog, Pou5f3, and Sox19b function independently and cooperatively to remodel the chromatin. (A) Heatmap showing the rescue in accessibility for all the regions that lost accessibility in MZ*nps* compared to wild-type (WT) (Group 1). Here, the rescue level is measured as the ratio between the accessibility in each condition and the wild-type (WT) accessibility for that region. Warmer color represents higher level of rescue. *S*-*high and S* represent a high and low Sox19b rescue dosage respectively (See Methods). (B, C) Bar plots comparing the number of regions in Group 1 (B) or in Group 1 co-bound by (C) that can be rescued in different conditions. Nanog shows the strongest pioneering activity. Rescuing with multiple factors simultaneously can rescue more regions than each single factor rescues. Here, an accessible region was defined as being rescued if a peak was called from Omni-ATAC data in that condition (at the significance level at *P* = 10^−8^) and overlap with the accessible region in WT. (D-F) Representative genome tracks showing the level of rescue in accessibility for Group 1 regions that is bound by all three factors in different conditions. (D) Regions that were rescued by only one of the three factors; (E) A region that can be rescued by all three factors individually; (F) A region that requires both Nanog and Pou5f3 to rescue, suggesting NPS function independently and cooperatively to establish chromatin accessibility. (G) Violin plot comparing the nucleosome occupancy between regions that can be rescued by a single, double or triple factor combination. Regions that require multiple factors to rescue have significantly lower nucleosome occupancy than regions that can be rescued by a single factor (N>NP: *P* = 3.4×10^−33^; N>NS: *P* = 1.0×10^−16^; P>NP: *P* = 5.0×10^−3^; P>PS: *P* = 1.1×10^−2^; S>NS: *P* = 2.9×10^−9^; S>PS: *P* = 2.7×10^−10^; NP>NPS: *P* = 8.6×10^−11^; NS>NPS: *P* = 1.6×10^−10^; PS>NPS: *P* = 5.3×10^−8^, two-sample t-test, * *P* <0.05,** *P* <0.01, *** *P* <0.001.). On the x-axis, ^†^ denotes regions that require two or all three factors for rescue, e.g., NP^†^ denotes regions that require both Nanog and Pou5f3 to rescue accessibility. (H) Stacked bar plot showing the percentage of Group 1 regions that require one, two, or three NPS factors to rescue accessibility. All regions were ranked according to the average nucleosome occupancy score within the region and were then divided into quartiles. Regions with lower nucleosome occupancy more frequently require multiple factors to rescue accessibility compared to those with higher nucleosome occupancy. (I) Violin plot comparing the nucleosome occupancy between regions that can be rescued by Sox19b at the low and high dosage. Regions that require the high dosage of Sox19b have significantly lower nucleosome occupancy than regions that can be rescued by the low dosage of Sox19b (*P* = 3.7×10^−10^, Two-sample t-test). S-high^†^ represents regions that can only be rescued by the high dosage of Sox19b.

To investigate synergy and/or redundancy in the pioneer-like activity of NPS, we focused on 1,613 regions that lost accessibility in MZ*nps* embryos (Group 1) and were co-bound by all three factors (Figures 4C and S4B). There were 8, 60, and 517 regions that are specifically accessible in MZ*nps* embryos when only Sox19b, Pou5f3, or Nanog function was restored, respectively. ChIP-seq analysis revealed that these regions were bound by all three factors in wild-type embryos, yet accessibility was restored by only one of the factors (Figures 4D and S4B). In contrast, in a subset of regions, two or more factors worked interchangeably to establish chromatin accessibility (Figure 4E). Interestingly, 801 regions only became accessible when simultaneously rescued by two or three factors, but not by a single factor (Figures 4B and 4F). These results demonstrate that Nanog, Sox19b and Pou5f3 can bind closed chromatin either alone, interchangeably, or synergistically to establish chromatin accessibility. Taken together these results indicate that the pioneering activity of Nanog, Pou5f3, and Sox19b is not purely an intrinsic property of the protein but is instead modulated by the sequence context and co-factors *in vivo*.

### Identification of sequence features determining the pioneering activity of Nanog, Pou5f3, and Sox19b

So far, our results demonstrate that binding of Nanog, Pou5f3 and Sox19b either alone or in combination with each other can initiate chromatin opening in certain contexts. However, the regulatory rules that define each context are unknown. In order to define the genetic features that determine the capacity for Nanog, Pou5f3, and/or Sox19b to open chromatin, we investigated the following features: motif scores and motif numbers (Grant et al., 2011) for Nanog, Pou5f3, and Sox19b, as well as the nucleosome occupancy score (Kaplan et al., 2009) comparing regions requiring single factors (N, P, S), two factors (PS, NS, NP), and all three factors (NPS). We observed that the motif frequency and strength for each factor was higher in regions rescued by the cognate factor (Figures S4C-S4E). Regions rescued by two factors had a combined motif frequency that was higher than those regions rescued by a single factor (Figure S4C). On the other hand, the motif score was higher in regions rescued by a single factor than in regions only rescued by all three factors, suggesting that those regions requiring multiple factors tend to have weaker motifs (Figure S4D and S4E). Interestingly, we observed that regions requiring a single factor had higher nucleosome occupancy scores than those rescued by two or more factors (Figure 4G). Conversely, sites with higher nucleosome occupancy scores were more likely to require fewer factors (Figure 4H). Led by these data, we hypothesized that the nucleosome itself might promote the recognition/binding of single factors to the chromatin, whereas regions with lower nucleosome affinity might require multiple pioneer factors to initiate chromatin opening at those sites. Consistent with this model, sites with stronger synergistic effects exhibited lower nucleosome occupancy scores (Figure S4F), and regions rescued by a lower concentration of Sox19b had higher nucleosome occupancy scores (Figure 4I). In summary, sites with lower nucleosome occupancy, lower binding strength, or fewer binding motifs are more likely regulated by the co-occupancy of multiple pioneer factors. These results indicate that in addition to motif binding features, nucleosome occupancy also contributes to the pioneer-like activity of Nanog, Pou5f3, and Sox19b *in vivo*.

### Nanog, Pou5f3, and Sox19b regulate H3K27ac and H3K18ac levels at both enhancers and promoters

Our results indicate that NPS are essential for establishing chromatin accessibility primarily at enhancers and less so at promoters. However, MZ*nps* mutants fail to activate a large fraction of the zygotic genome. These observations raise an intriguing question: what is the mechanism by which NPS trigger transcriptional activation if it is not via promoter accessibility? This trigger should be specific to active genes, dependent on NPS, and coincide with transcriptional activation. Among different epigenetic landmarks, histone acetylation is often associated with chromatin opening and active transcription (Chan et al., 2019; Garcia-Ramirez et al., 1995; Hebbes et al., 1994; Krajewski and Becker, 1998; Nightingale et al., 1998; Tse et al., 1998). In *Drosophila*, the appearance H3K18ac, H3K27ac, and H4K8ac coincides with ZGA and correlates with the binding of the pioneer factor Zelda (Li et al., 2014), yet whether these changes are causal or simply correlate with transcription is not fully understood. In addition, histone acetylation is also associated with early zygotic transcription and premature expression of p300 and BRD4 can prematurely activate and increase the levels of zygotic transcription in zebrafish embryos (Chan et al., 2019). To determine how NPS remodel the epigenetic landscape to trigger the activation of the zygotic genome, we set out to determine how the loss of Nanog, Pou5f3, and Sox19b affects histone acetylation. Two lines of evidence suggest that NPS mediate ZGA through histone acetylation. First, we analyzed the relationship between H3K18ac, H3K27ac, and H4K8ac and the onset of genome activation. Similar to H3K27ac (Chan et al., 2019; Sato et al., 2019), the time course analysis of H4K8ac and H3K18ac by immunostaining revealed that these marks increase during ZGA (Figure S5A). However, only H3K27ac and H3K18ac are enriched at *miR-430* loci, one of the firstly transcribed loci, at the onset of zygotic transcription at 2.5 hpf (Figures 5A and S5B). *miR-430* transcription depends on NPS function as determined by Click-iT-seq and Click-iT imaging (Figures 1E and 5B). Interestingly, the enrichment of H3K27ac and H3K18ac at the *miR-430* locus was completely lost in MZ*nps* embryos (Figures 5A and S5B). Consistently, ChIP-seq signal for H3K27ac and H3K18ac was strongly reduced at the *miR-430* locus in MZ*nps* embryos (Figure S5C). These results suggest that H3K27ac and H3K18ac depend on NPS and that their deposition may mediate the activation of the first zygotically expressed gene *miR-430*.

**Figure 5.**
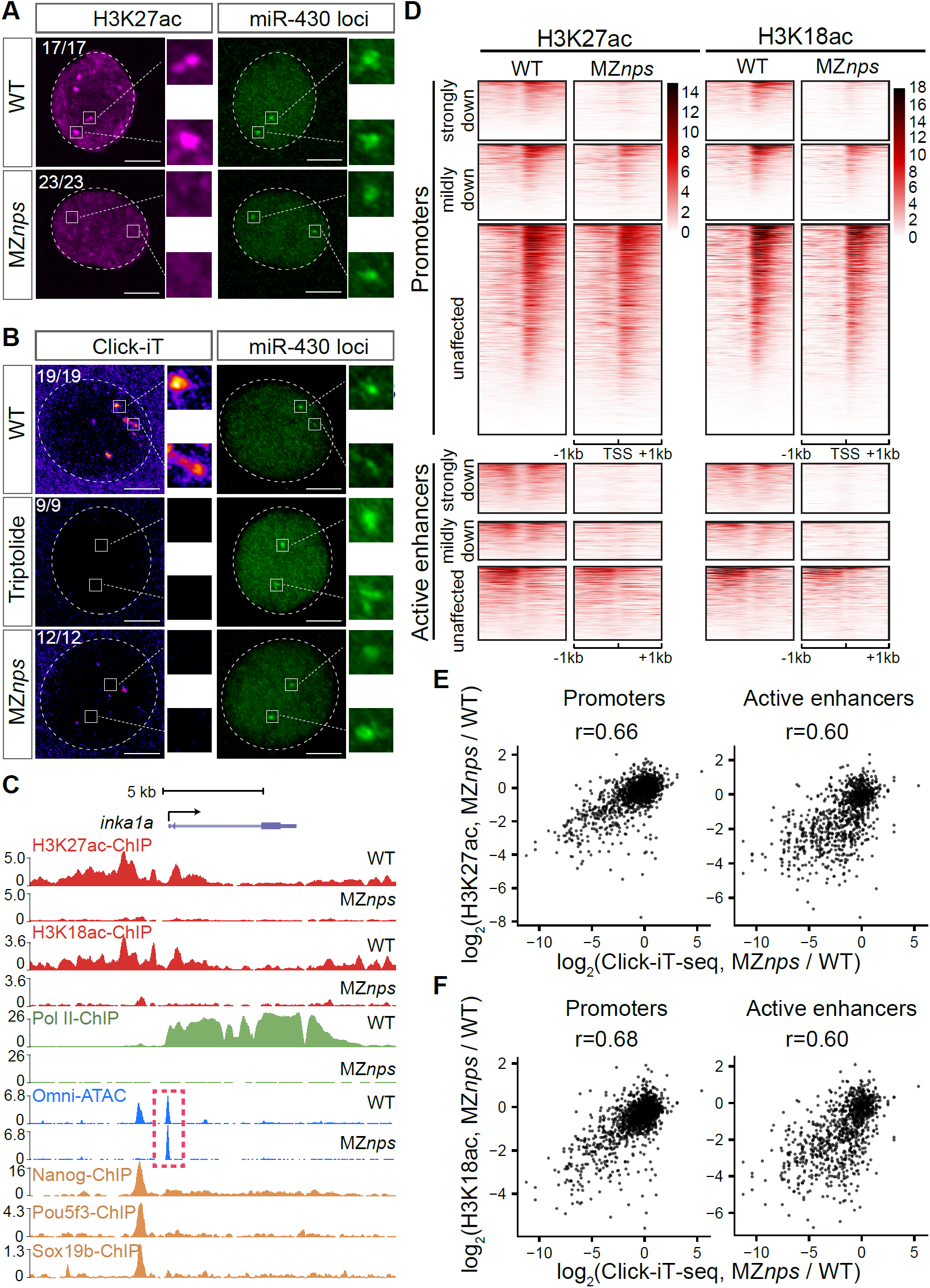
Loss of Nanog, Pou5f3, and Sox19b leads to loss of H3K27ac and H3K18ac at promoters and enhancers. (A) Confocal imaging of immunofluorescence (IF) against H3K27ac at 2.5 hpf, with labeling of the *miR*-*430* locus with Cas9-3xGFP. In wild-type (WT) embryos, H3K27ac was enriched at the *miR*-*430* locus, but is lost in MZ*nps* embryos. Scale bar: 5 µm. Nuclei are outlined with a dashed line based on the DAPI signal. (B) Confocal imaging labeling nascent transcription by Click-iT imaging and the *miR-430* locus with Cas9-3xGFP at 2.5 hpf. In WT embryos, the *miR*-*430* locus colocalized with nascent transcripts. Triptolide blocks transcription of all loci, including *miR*-*430*. In MZ*nps* embryos, some loci were transcriptionally active, but no transcription was detected at the *miR*-*430* locus. Scale bar: 5 µm. (C) Representative genomic tracks for H3K27ac, H3K18ac, Pol II binding, accessibility, and NPS binding intensities for the gene *inka1a*, whose transcription is strongly downregulated in MZ*nps*. In MZ*nps* embryos, the promoter of *inka1a* remained accessible (dash line), but H3K27ac and H3K18ac were dramatically decreased at both the promoter and nearby enhancers, and Pol II was also largely reduced. (D) Heatmaps showing H3K27ac and H3K18ac levels at promoter and active enhancer regions of different categories of zygotic genes in WT and MZ*nps* embryos. Genes that were strongly downregulated in MZnps exhibited the most dramatic reduction in H3K27ac and H3K18ac levels in MZ*nps* embryos at both promoters and active enhancers. Heatmaps are centered at the TSS and the summit of active enhancers with 500 bp on both sides and are ranked by the average H3K27ac signal within the flanking 1000 bp region from high to low. (E, F) Biplot showing high correlation between change in H3K27ac level (E) or H3K18ac level (F) in (MZ*nps*/WT) and the change in transcription level (MZ*nps*/WT) at active enhancers and promoters (r, Pearson correlation).

Second, to analyze the genome-wide effect of NPS loss-of-function on H3K27ac and H3K18ac, we performed ChIP-seq for H3K27ac and H3K18ac in MZ*nps* and wild-type embryos at 4 hpf. We observed a major reduction in H3K27ac and H3K18ac at both enhancers and promoters for the majority of the downregulated genes (Figures 5C, 5D and S5C-S5E). For example, strongly downregulated genes such as *inka1a* and *klf17* lost H3K27ac and H3K18ac in MZ*nps* embryos at both promoters and enhancers (Figures 5C and S5C). In contrast, most unaffected genes showed similar H3K27ac and H3K18ac levels at enhancers and promoters between wild-type and MZ*nps* embryos (i.e. *actb1*) (Figures 5D and S5C-S5E). Additionally, the reduction in H3K27ac and H3K18ac correlated with the reduction in transcription in MZ*nps* embryos (Figures 5E and 5F), suggesting that the lack of H3K27ac and H3K18ac are associated with the loss of transcriptional activation in MZ*nps* embryos.

We observed a stark contrast between chromatin accessibility and histone acetylation (H3K27ac and H3K18ac) at enhancers and promoters. Loss of NPS reduced enhancer accessibility but promoter accessibility was largely unaffected, consistent with the stronger binding of NPS at enhancers (Figure 2B). In contrast, loss of NPS reduced H3K27ac and H3K18ac at both enhancers and promoters for strong target genes, such as *inka1a* (Figures 5C and 5D). Thus, the TFs maintaining accessibility at promoters are not sufficient to recruit histone acetyltransferases or mediate Pol II recruitment and transcriptional activation in the absence of NPS. Taken together, these results indicate that NPS are required for the recruitment of histone acetyltransferases and the acetylation of H3K27, H3K18 and possibly other histone residues to mediate genome activation.

### Histone acetylation but not chromatin opening can bypass the function of Nanog, Pou5f3, and Sox19b to activate target genes

Our results suggest that NPS reprogram the chromatin for ZGA by recruiting histone acetyltransferases and subsequent histone acetylation. If this is true, then we would expect the recruitment of histone acetyltransferases and restoration of histone acetylation in MZ*nps* embryos to rescue the transcription of NPS target genes (Figure 6A). To test this, we attempted to rescue histone acetylation by specifically directing histone acetyltransferase activity to two strong NPS target genes, *asb11* and *her5*. We fused the histone acetyltransferase core domain (HAT) of human p300 (Hilton et al., 2015) to a catalytically dead Cas9 (Chan et al., 2019) (dCas9-HAT) and used four sgRNAs to target this complex to the regulatory regions of *asb11* and *her5*. We found that dCas9-HAT targeting *asb11* or *her5* activated their transcription in MZ*nps* embryos as shown by qPCR and RNA *in situ* hybridization. As a control, we observed that activation of these genes was dependent on the specific sgRNAs targeting that locus, indicating that the rescue was specific to genes targeted (Figure 6B). We also demonstrated that dCas9 occupancy itself cannot restore transcription independent of the histone acetyltransferase activity by fusing a catalytically dead HAT to dCas9 (dCas9-dHAT), where dHAT contains an inactivating mutation (D1399Y) in the HAT (Hilton et al., 2015) (Figures 6B and S6A). Although the dCas9-HAT rescue of transcription did not reach wild-type levels (Figure 6B), this is likely due to the mosaic nature of the rescue caused by microinjection. Indeed, RNA *in situ* hybridization for *asb11* and *her5* revealed that both genes are transcribed at high levels in large clonal patterns (Figure 6C).

**Figure 6.**
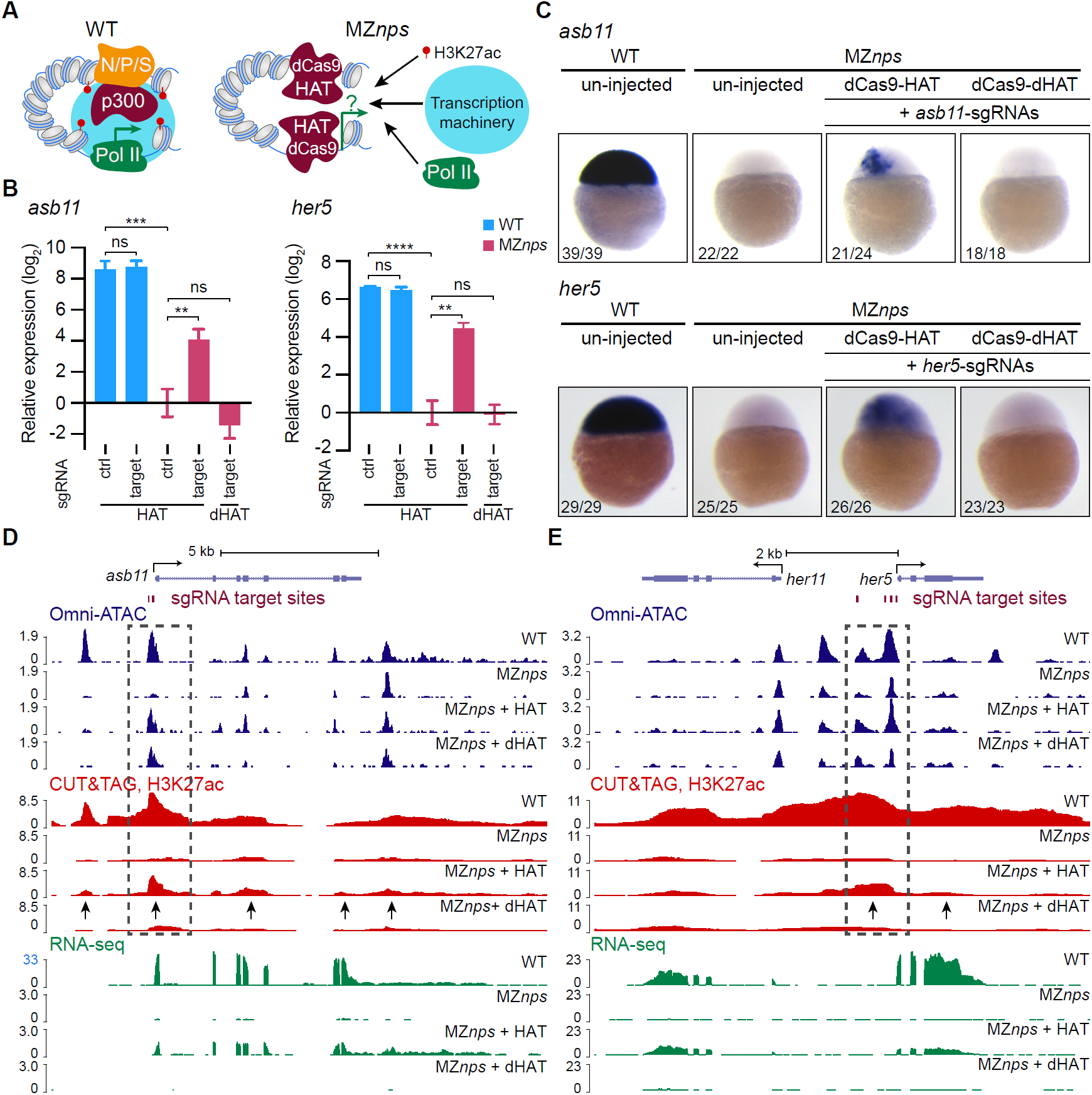
Restoration of H3K27ac partially rescues transcription of Nanog, Pou5f3, and Sox19b strong targets. (A) Schematic illustrating the experiment to test whether recruiting histone acetyltransferase activity can bypass the function of NPS in activating transcription. The dCas9-HAT system was used to introduce histone acetylation at target sites. (B) qPCR revealed that transcription of *asb11* and *her5* was partially restored by dCas9-HAT targeting the regulatory regions of *asb11* and *her5*, respectively, in MZ*nps* embryos. dCas9-HAT together with sgRNAs targeting *asb11*, but not control sgRNAs, activated *asb11*; and Cas9 with catalytically dead HAT dCas9-dHAT failed to activate *asb11*. Similarly, only dCas9-HAT together with sgRNAs targeting *her5* activated *her5*. (C) RNA *in situ* hybridization was used to detect the expression pattern of *asb11* and *her5* in wild-type (WT) and MZ*nps* embryos. dCas9-HAT system restored transcription in MZnps mutant embryos. (D) Genome tracks showing that accessibility and H3K27ac were restored at the *asb11* promoter region, and transcription of *asb11* was also restored in MZ*nps* embryos by the dCas9-HAT system. Signal intensity in RPM (reads per million). (E) Genome tracks showing that H3K27ac was restored at the targeted *her5* promoter region and the enhancer region near the TSS of *her5*. Transcription was activated for both *her5* and *her11*, a gene near *her5*. Signal intensity in RPM. The location of the gRNAs used in each experiment are indicated in the track with red bars and the dashed line box. Arrows show regions were H3K27Ac is rescued.

To determine whether rescue by dCas9-HAT was mediated by histone acetylation rather than chromatin opening at the promoter caused by the dCas9 targeting itself, we performed RNA-seq, Omni-ATAC, and H3K27ac CUT&TAG in MZ*nps* embryos injected with dCas9-HAT targeting both *asb11* and *her5* simultaneously. We observed that in MZ*nps* embryos, dCas9-HAT partially restored both H3K27ac and transcription of *asb11* and *her5*, but had no effect on a control gene that was not targeted by these sgRNAs (*klf17*) (Figures 6D, 6E, and S6B). In contrast, dCas9-dHAT restored chromatin accessibility at the *asb11* promoter, but not H3K27ac or transcription of *asb11* in MZ*nps* embryos (Figure 6D), further suggesting that opening of the promoter or the enhancer is insufficient to promote transcription during ZGA. Interest-ingly, targeting dCas9-HAT to the regulatory regions up-stream of the *her5* locus also restored transcription of the neighboring gene *her11*(Figure 6E), suggesting a causal relationship by which a local increase in histone acetylation is sufficient to recruit Pol II and restore transcription at both promoters, independently of NPS (Figure S6C). It is worth noting that targeting dCas9-HAT to the promoter of *asb11* was also able to increase H3K27ac at upstream and down-stream regions above the levels observed with dHAT. These sites are also found to be acetylated in wild-type embryos, and this observation offers the possibility that the local conformation of the chromatin likely brings those regions in close proximity to the promoter where dCas9-HAT can mediate the acetylation of H3K27, even in the absence of NPS. Altogether, these results demonstrate that recruiting histone acetyltransferase activity with depositing H3K27ac in enhancer and promoter regions is sufficient to activate zygotic genes independent of the pioneer function of NPS, indicating that regulation of histone acetylation is a critical step through which Nanog, Pou5f3 and Sox19b regulate activation of the zygotic genome.

## Discussion

This study provides three major insights into the mechanisms that initiate the awakening of the zygotic genome by identifying the network of TFs that regulate chromatin accessibility together with Nanog, Pou5f3, and Sox19, defining how genomic sequence context modulates the pioneering function of NPS *in vivo*, and elucidating the causal relationship between the binding of these factors and transcripttion via epigenetic remodeling through histone acetylation.

First, we identified co-occupancy networks of maternally expressed TFs whose DNA binding motifs overlap with accessible chromatin during genome activation and defined their dependency on NPS. Recent studies have investigated the role of individual transcription factors in embryonic stem (ES) cells and in early embryos (Charney et al., 2017; Gao et al., 2018; Gao et al., 2020; Lu et al., 2016; Pálfy et al., 2020; Schulz et al., 2015; Soufi et al., 2015; Sun et al., 2015; Veil et al., 2019b), providing evidence that individual factors can prime a limited fraction of the genome. However, analysis of single mutants is limited by the fact that >50% of NPS-bound accessible sites are bound by at least two of these factors, suggesting a combinatorial role of NPS in regulating ZGA in zebrafish (Lee et al., 2013). Given the functional redundancy of these TFs at many genomic sites, any single mutant will provide an underestimate of that factor’s regulatory activity as the loss-of-function can be compensated for by the other two factors. In contrast to previous studies, we found that NPS are responsible for chromatin opening at more than half of active enhancers during the MZT, and that loss of enhancer accessibility is correlated with a reduction in transcription. These results demonstrate that NPS play critical roles in establishing enhancer accessibility during ZGA. In contrast, NPS appear to be dispensable for promoter opening at most NPS-dependent genes. While Pol II is responsible for promoter opening in *Drosophila* (Gilchrist et al., 2008; Lee et al., 1992; Levine, 2011; Shopland et al., 1995), our ChIP-seq data in MZ*nps* embryos revealed that Pol II recruitment is largely reduced in the absence of NPS, indicating that other TFs, rather than Pol II, are responsible for promoter opening in MZ*nps* embryos. These results suggest that the establishment of chromatin accessibility at promoters and enhancers are independent processes during the MZT, where the pioneering activity of NPS contributes to, but is not obligatory for the opening of most promoters. While the loss of NPS leads to a dramatic reduction in accessibility at many of NPS binding sites, a significant fraction of NPS binding sites remain accessible in MZ*nps* embryos, indicating that other TFs co-occupy the same regulatory regions and can initiate chromatin opening independently of NPS. Our analysis of TF expression and motifs enriched in accessible chromatin revealed an extensive network of TFs associated with chromatin accessibility during ZGA. For example, we found the SP1, NFY, FOXK1, BBX, and KLF9 binding motifs to be enriched at promoters and enhancers, suggesting that these factors may orchestrate genome activation independently of the pioneering activity of NPS. Indeed, NFY and FoxH1 regulate early zygotic gene expression in mouse during preimplantation development and in *Xenopus* during mesendoderm formation (Charney et al., 2017; Lu et al., 2016). In contrast, EOMES motif was enriched at sites where accessibility depends on NPS, and we demonstrated that Eomesa binding to chromatin is dependent on the pioneering activity of NPS for the majority of its target regions. Similarly, in *Drosophila*, Zelda regulates chromatin binding for Bicoid and Dorsal (Schulz et al., 2015; Sun et al., 2015). These results indicate that, in addition to activating their target genes, TFs such as Nanog, Pou5f3, Sox19b and Zelda also function to pioneer the opening of the chromatin for other TFs during ZGA to reprogram the silent chromatin and orchestrate gastrulation and development. While most TFs that function independently of NPS are responsible for housekeeping functions, TFs that depend on NPS (e.g., Eomesa) and NPS target genes are enriched in developmental functions. These results suggest that the fertilized egg is re-programmed by different cohorts of TFs that regulate house-keeping programs and developmental programs (Eisenberg and Levanon, 2013; Lam et al., 2012; Murphy et al., 2018; Rach et al., 2011; Zabidi et al., 2015), with NPS being the major drivers of developmental programs in zebrafish. Consistently, mammalian NANOG, OCT4, and SOX2 co-occupy many genomic loci and are required for pluripotency and lineage commitment (Boyer et al., 2005; Chen et al., 2008; Loh et al., 2006; Wang et al., 2012). Altogether, by uncovering TFs associated with chromatin accessibility, their motif co-occurrence, and their dependency on NPS, our results have revealed a complex TF network underlying developmental reprogramming during the MZT.

Second, the interplay between Nanog, Pou5f3, and Sox19b in establishing chromatin accessibility depends on the genomic context and nucleosome occupancy. We found that each single factor displays pioneering activity *in vivo* and can initiate chromatin opening in the absence of the other two factors. We also observed that Nanog, Pou5f3, and Sox19b can function interchangeably to initiate the opening of the chromatin at many loci. Due to this functional redundancy, the pioneering activity of NPS can easily be under-estimated by analyzing individual mutants (Gao et al., 2020; Pálfy et al., 2020; Veil et al., 2019a). We hypothesize that redundancy in pioneering activity may provide another level of regulation, and, together with shadow/redundant enhancers (Cannavò et al., 2016; Hong et al., 2008), may be essential for robustness of transcriptional regulation of key developmental genes. In contrast, other regions are pioneered by a combination of NPS, where these factors work cooperatively or additively to open the chromatin because no individual factor is able to open the chromatin independently. This scenario might provide a higher level of spatial or temporal precision in gene regulation by integrating multiple inputs. These sites are characterized by higher NPS motif co-occurrence, weaker NPS motifs (lower similarity to the consensus motifs) (Meers et al., 2019), and lower nucleosome occupancy scores. Nucleosomes often inhibit TF binding; however, many pioneer/pioneer-like factors show preferential binding at high intrinsic nucleosome sites (Iwa-fuchi-Doi and Zaret, 2014, 2016; Zaret and Carroll, 2011; Zhu et al., 2018). For example, the pioneer factors OCT4 and SOX2 preferentially bind sites enriched for nucleosomes to initiate reprogramming of human fibroblasts (Soufi et al., 2015). Similarly, binding sites for NPS in zebrafish embryos, Zelda in *Drosophila* embryos, and progesterone receptor in breast cancer cells are enriched for nucleosome occupancy (Ballaré et al., 2013; Gao et al., 2020; Schulz et al., 2015; Sun et al., 2015; Veil et al., 2019b). It is generally believed that these pioneer/pioneer-like factors must overcome the nucleosome barrier to facilitate the binding of other TFs and to activate transcription (Pálfy et al., 2020; Zaret and Carroll, 2011). However, our results suggest that nucleosomes facilitate the pioneering activity of NPS in vivo. This may be caused by the differential affinity of the TFs for their cognate binding sites depending on the relative position with respect to the nucleosome (Li et al., 2019; Michael et al., 2020; Zhu et al., 2018). Indeed, we observed that sites that become accessible at lower TF concentrations or that are pioneered by a single factor have higher nucleosome occupancy scores. Additionally, Eomesa binding sites that are independent of NPS have higher nucleosome occupancy that the dependent ones, suggesting that the nucleosome might also facilitate Eomesa binding to chromatin. These results are consistent with previous studies in human, where EOMES is able to bind nucleosomal DNA and has been suggested to have pioneering activity during differentiation (Meers et al., 2019; Zhu et al., 2018). Altogether, our results support that nucleosomes facilitate pioneering activity and provide a deeper understanding of how NPS interplay with each other to achieve developmental reprogramming during the MZT.

Third, we demonstrated that the recruitment of histone acetyltransferase (p300) can bypass the function of NPS to control ZGA. Conversely, we observed a reduction of histone acetylation (H3K27ac and H3K18ac) in MZ*nps* embryos across enhancers and promoters of NPS target genes. These results suggest that NPS mediate ZGA through regulating histone acetylation. Studies in ES cells have shown that BRD4 recognizes acetylated histones (Filippakopoulos et al., 2012; Shi and Vakoc, 2014) and promotes transcriptional initiation and elongation by recruiting the Mediator complex and p-TEFB (Cho et al., 2018; Gibson et al., 2019; Itzen et al., 2014; Sabari et al., 2018; Yang et al., 2008). Our analysis of Pol II ChIP-seq revealed that NPS primarily function in recruiting, rather than releasing, Pol II. Based on these results, we propose a model where NPS activate the zygotic genome by recruiting Pol II, possibly through Brd4 and the Mediator complex, in a process that is initiated by histone acetylation. Consistent with this model, our previous studies have shown that the function of p300 and Brd4 is required for ZGA (Chan et al., 2019), and Med13 is required for ZGA in mice (Miao et al., 2018). BRD4 is required for the clustering of Mediator and Pol II at super-enhancers (Sabari et al., 2018), which are associated with high levels of pluripotency factors (e.g., NANOG, OCT4, and SOX2), Mediator, and H3k27ac in ES cells (Hnisz et al., 2013; Whyte et al., 2013). Interestingly, we observed that H3K27ac and H3K18ac at the *miR*-*430* locus depend on NPS and appear to form puncta, as does P-ser5 Pol II (Chan et al., 2019; Hadzhiev et al., 2019; Sato et al., 2019). Histone acetylation is able to dissolve chromatin droplets and the binding of BRD4 promotes a new liquid phase of the acetylated chromatin *in vitro* (Gibson et al., 2019). Future studies will be needed to understand whether NPS regulate genome activation *in vivo* through liquid-liquid phase separation and how these changes affect chromatin conformation of individual loci. It has been suggested that histone acetylation facilitates chromatin opening (Garcia-Ramirez et al., 1995; Hebbes et al., 1994; Krajewski and Becker, 1998; Tse et al., 1998). We observed that loss of H3K27ac and K18ac at promoters in MZ*nps* embryos did not lead to a loss of accessibility at the promoters, indicating that H3K27ac and K18ac are dispensable for chromatin accessibility during ZGA. Finally, we found that NPS can regulate H3K27ac at promoters distant from their main binding site at the enhancer. Conversely, we find that dCas9-HAT binding at promoters can acetylate nearby enhancers, even in the absence of NPS. Our results suggest that 3D proximity between enhancers and promoters facilitates the acetylation of both when histone acetyltransferase is recruited to either region (Figure 6D). Interestingly, we observed that this structural bias at the tested loci is maintained even in the absence of NPS, suggesting that this is either the intrinsic chromatin organization or that other factors are mediating promoter-enhancer interactions at these loci. Future studies will be needed to investigate the causal relationship between NPS binding and the 3D organization of the chromatin during ZGA. Altogether, our results suggest that NPS potentiate the genome by initiating chromatin opening and activate the zygotic genome by regulating histone acetylation, providing novel insights to the molecular mechanisms underlying genome activation and reprogramming during the MZT.

## Acknowledgments

We thank Steven Henikoff for sharing pA-Tn5 protein and advice on CUT&TAG; Fiona Wardle for sharing the Eomesa antibody; Jose Luis Gomez-Skarmeta and Juan Ramon Martinez Morales for sharing their protocols and ATAC-seq data in zebrafish. Karen Adelman and Bluma Lesch for mentorship in ChIP-seq; Yong Zhang and Guifen Liu for advice on ATAC-seq. We thank Kaya Bilguvar and Christopher Castaldi from the Yale Center for Genome Analysis for sequencing support; Hiba Codore, Nitya Khatri, Eliakim de Guzman, Sarah Dube, Valeria Schmidt, and Timothy Gerson for technical help. We thank Caroline Hendry, Bluma Lesch, and Maria Benitez for feedback on the manuscript. We thank the Surdna Foundation and the Yale Genetics Venture Fund for supporting L.M.; The Jane Coffin Childs Memorial Fund for supporting M.L.K.; the 2T32HD007149-41A1 NIH training grant for supporting M.E.P. This work is supported by NIH grants R01 HD074078, GM103789, GM102251, GM101108, and GM081602, R35 GM122580 and the 4DNucleome program.

## Author Contributions

L.M. and A.J.G. conceived the project and wrote the manuscript, with contributions from all authors. L.M. performed experiments, with contribution from A.R.B. and M.L.K. in ChIP-seq; and S.H.C. and M.E.P. in immunofluorescence imaging. Y.T. performed the computational analysis with contributions from C.E.V.; L.M., Y.T. and A.J.G. performed data analysis and interpreted the results with contribution from other authors. A.J.G. supervised the project.

---

**Figure S1.**
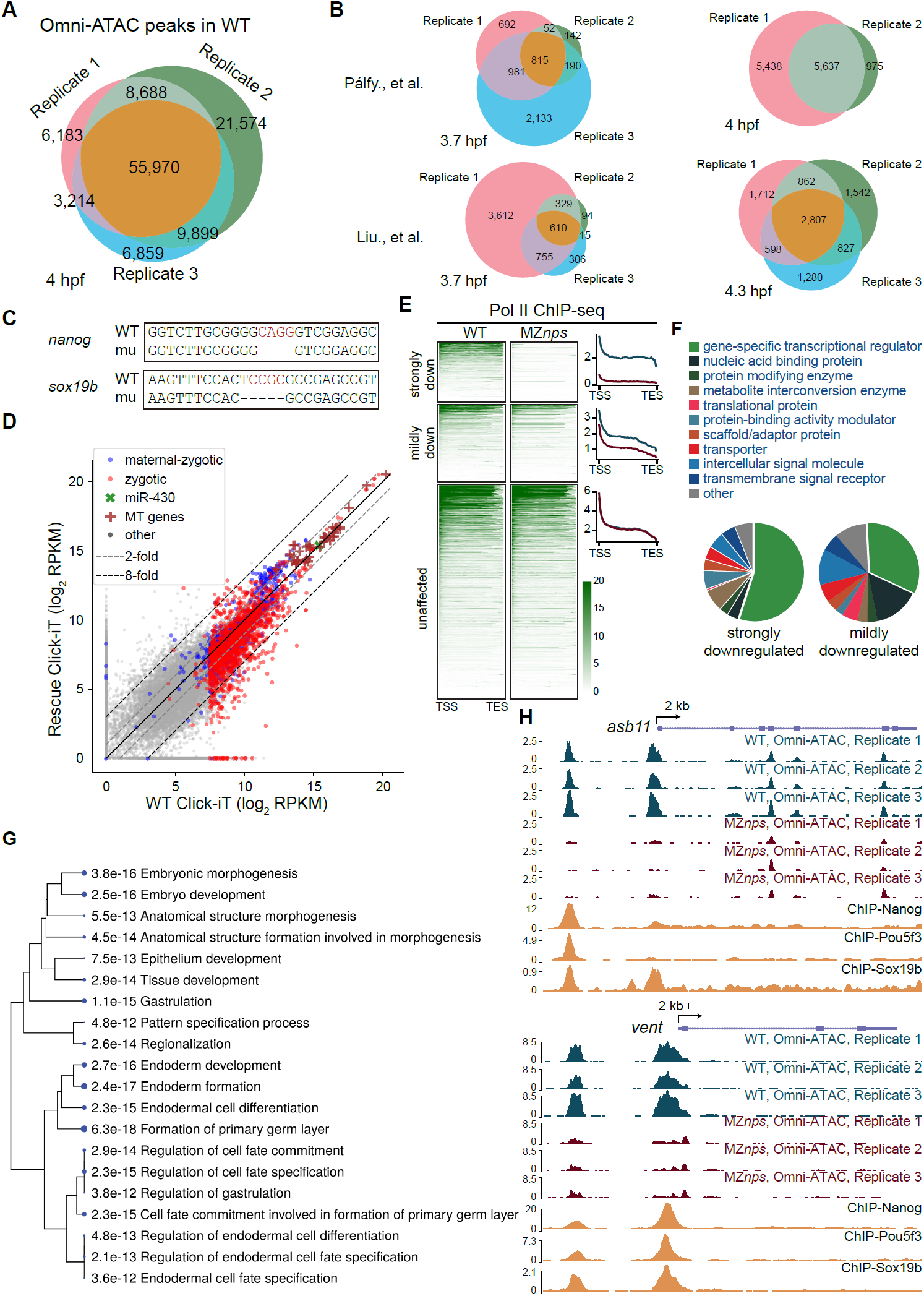
Loss of Nanog, Pou5f3, and Sox19b decreases chromatin accessibility. (A) Venn diagram showing high reproducibility across three replicates of Omni-ATAC replicates in wild-type (WT) embryos at 4 hpf. (B) Venn diagram showing the overlap of previously published ATAC-seq replicates in WT at 3.7-4.3 hpf. (C) DNA sequences of the *nanog* mutant (mu) and *sox19b* mutant generated with the CRISPR/Cas9 system. (D) Biplots comparing the nascent transcriptome of MZ*nps* embryos rescued with *nanog* and *pou5f3* mRNAs (Rescue) with that of WT embryos. (E) Heatmaps and line plots showing Pol II binding along the gene body of different categories of zygotic genes according to their level of downregulation in MZ*nps* compared to WT. Each zygotic gene was grouped into 100 bins from the TSS to the TES (transcription end site, annotated from Ensembl) and Pol II signal was averaged within each bin. (F) Protein class enrichment of the strongly downregulated and mildly downregulated genes as determined by PANTHER gene ontology enrichment analysis. (G) Hierarchical clustering tree showing the correlation among the top 20 significantly enriched biological processes for the strongly downregulated genes. The analysis was performed using ShinyGO gene ontology enrichment analysis. Biological processes are clustered together based on shared genes. Dot size indicates *P*-value: a larger size represents a lower *P*-value. (H) Representative genomic tracks of Omni-ATAC replicates in WT and MZ*nps* embryos, and NPS ChIP-seq.

**Figure S2.**
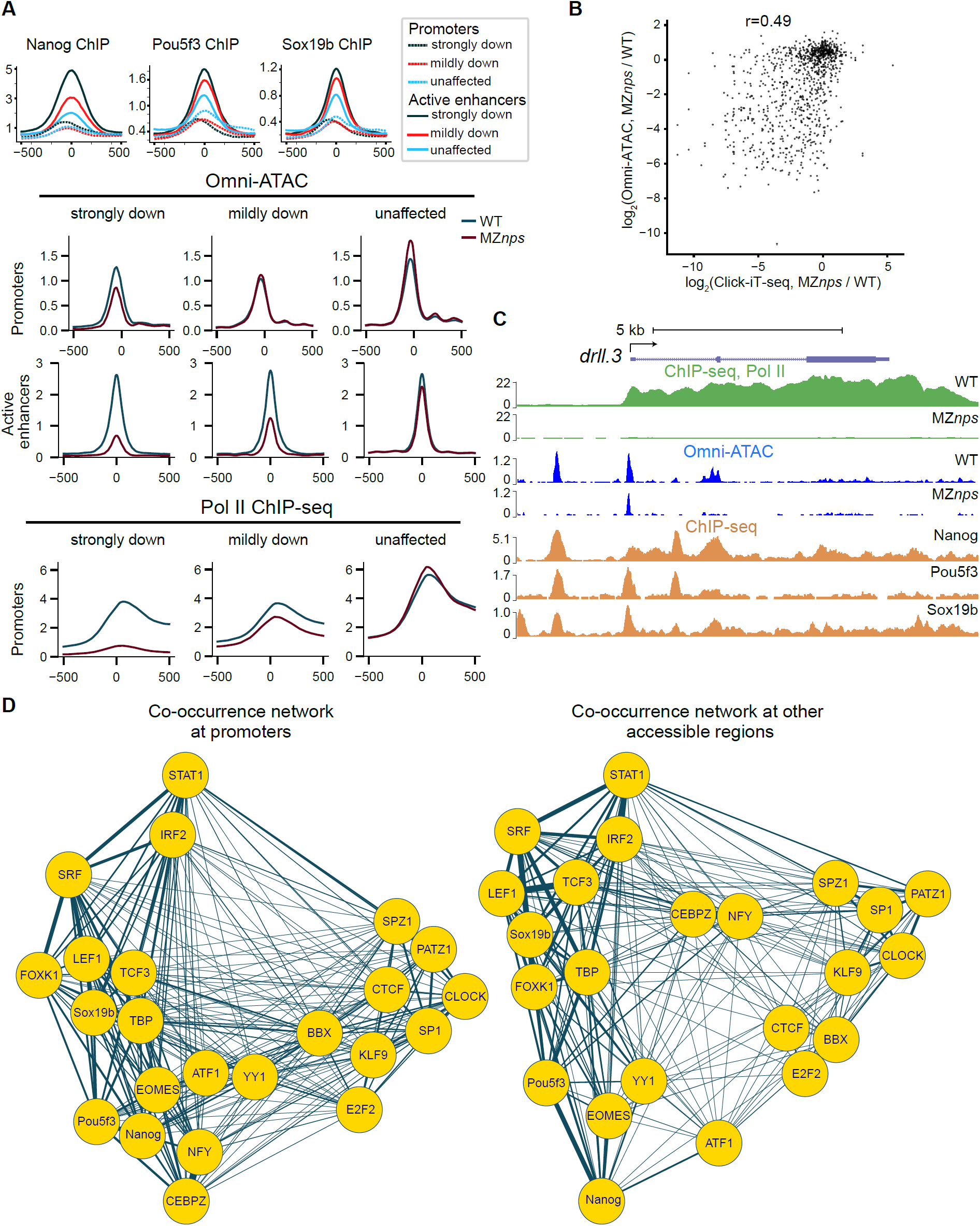
Accessibility at promoters is not established by Pol II. (A) Line plots comparing Pol II binding (bottom), accessibility (middle), and NPS binding intensities (top) at promoters and active enhancers between different groups of zygotic genes. (B) Biplot showing a high correlation between the change of Omni-ATAC and change of transcription in (MZ*nps*/WT) (r = 0.49, Pearson correlation) at active enhancers. (C) Representative genomic tracks of a *strongly downregulated* gene *drll*.*3*, showing that accessibility at the promoter is reduced but maintained, despite the loss of Pol II, suggesting that the remaining accessibility is not established by Pol II. (D) Co-occurrence of TF motifs at promoters and other accessible regions in Figure 2C. The closer distance and thicker connection between nodes represent higher co-occurrence between the sequence motifs of the TFs in the network.

**Figure S3.**
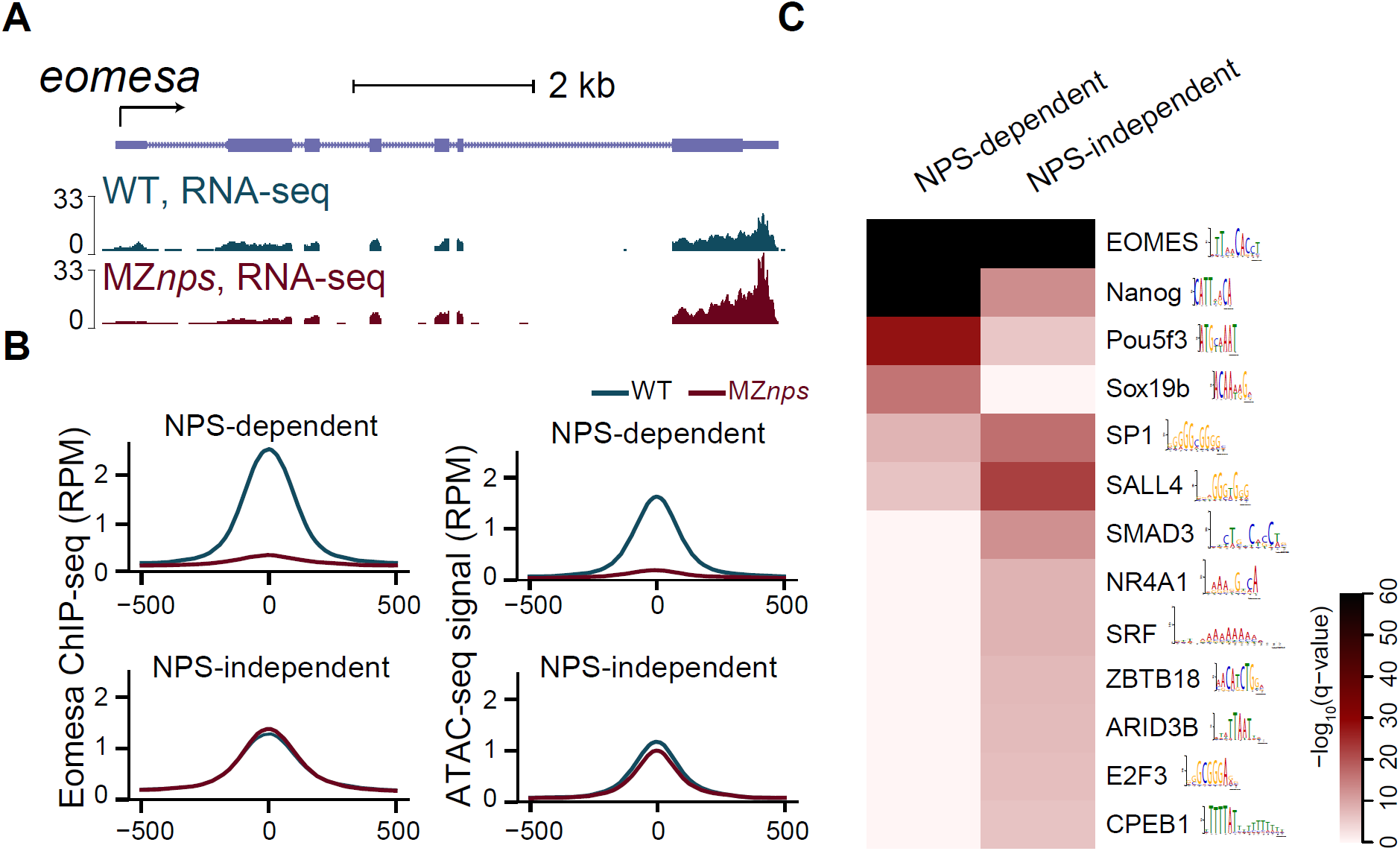
MZ*nps* embryos exhibit reduced Eomesa binding, but no reduced Eomesa transcript levels. (A) Genomic tracks of RNA-seq showing that *eomesa* expression in wild-type (WT) and MZ*nps* embryos. (B) Line plots showing the average Eomesa binding signal and accessibility at NPS-dependent and NPS-independent Eomesa binding regions in WT and MZ*nps* embryos. (C) Enrichment of known TF motifs at NPS-dependent and NPS-independent Eomesa binding regions. A random selection of 754 regions from both groups were taken to make the significance comparable between groups.

**Figure S4.**
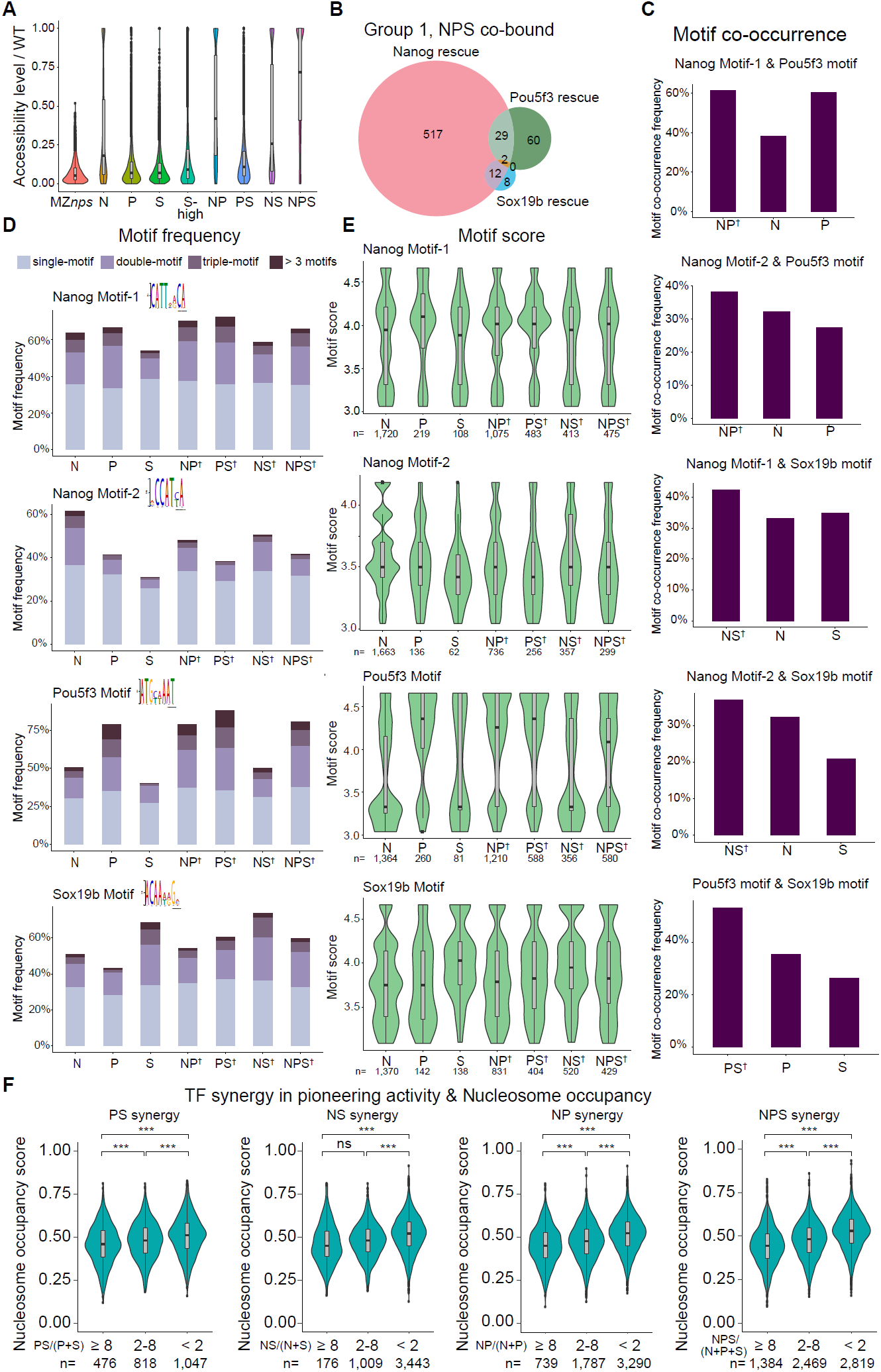
Motif frequency and strength determine the genomic context for Nanog, Pou5f3, and Sox19b pioneering activity. (A) Violin plot comparing the rescue level (ratio of accessibility in a given condition to that in wild-type (WT) embryos) of all the Group 1 regions across different conditions. Nanog shows the strongest pioneering activity compared to Pou5f3 and Sox19b. (B) Venn diagram showing the overlap of Group 1 regions bound by all three factors (co-bound) that can be rescued with a single NPS factor, suggesting that the binding of the factor does not determine whether the region can be rescued by the specific factor. (C) Bar plots comparing motif co-occurrence between regions that can be rescued by a single factor and those that require two factors. Regions that require two factors have a higher frequency of motif co-occurrence from each of the factors compared to regions that can be rescued by a single factor. (D-E) Stacked bar plots and violin plots comparing the motif frequency (D) and motif strength (E) between regions that can be rescued by a single factor, two factors only (marked by ^†^), or three factors only (marked by ^†^). (F) Violin plot comparing the nucleosome occupancy of Group 1 regions with high (≥ 8-fold), medium (2 ≤fold change <8) and low (0.5 <fold change <2) synergistic effect between multiple factors. Synergistic effect was measured as the ratio of accessibility when rescued by multiple factors simultaneously to the sum of accessibility when those factors were applied separately. Regions with high synergic effect between multiple factors have significantly lower nucleosome occupancy compared to regions with low synergic effect (*** *P* <0.001; two-sample t-test).

**Figure S5.**
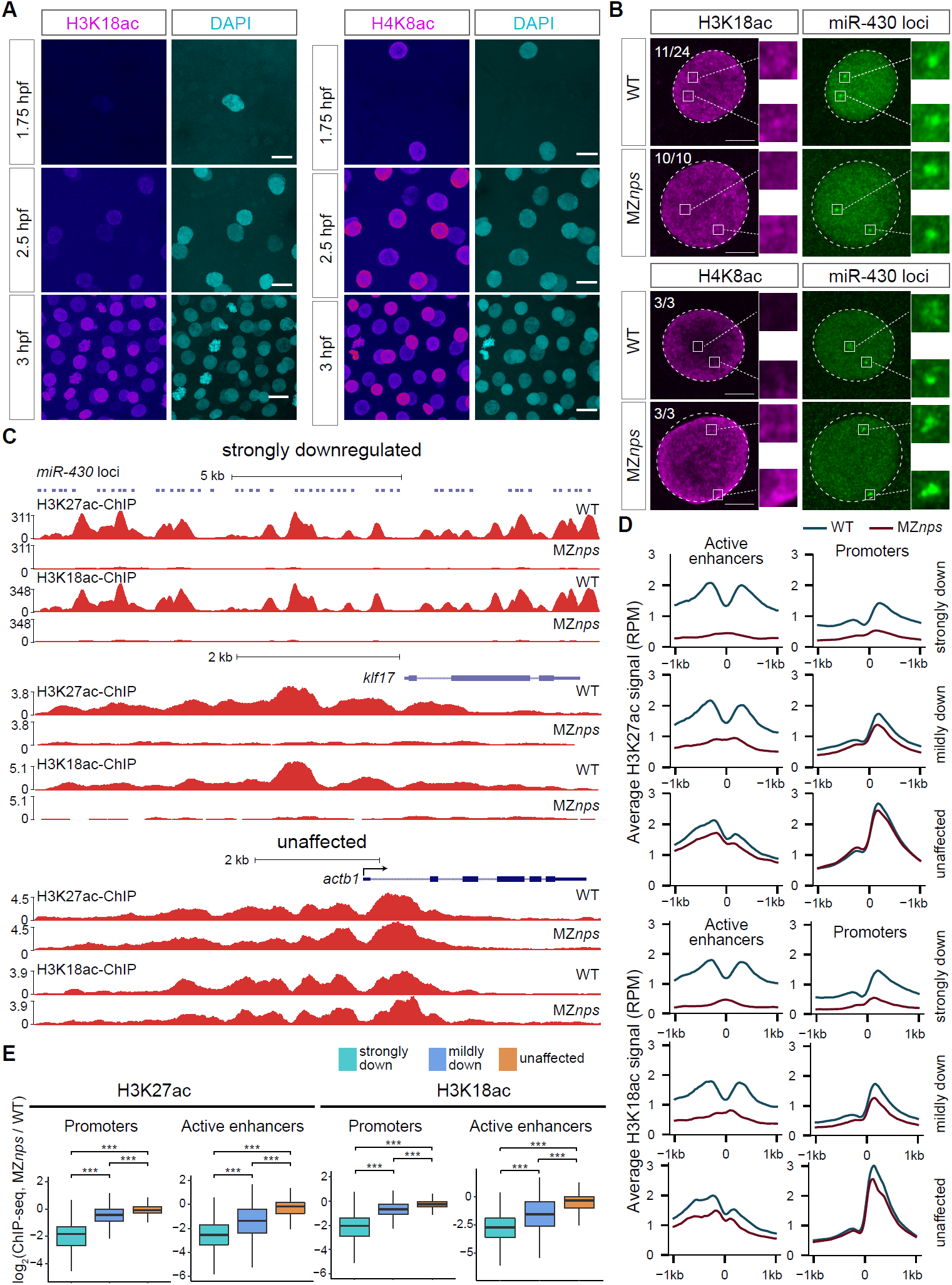
Loss of Nanog, Pou5f3, and Sox19b leads to a loss of H3K27ac and H3K18ac at promoters and enhancers of Nanog, Pou5f3, and Sox19b-dependent genes. (A, B) Confocal imaging of H3K18ac and H4K8ac immunofluorescence (IF) (red), DAPI (Blue) and the *miR*-*430* locus (Cas9-3xGFP, green). Scale bar in (A): 15 µm; Scale bar in (B): 5 µm. (C) Genomic tracks of H3K27ac and H3K18ac ChIP-seq showing H3K27ac and H3K18ac were largely reduced at the *miR*-*430* locus and *klf17* in MZ*nps* embryos. Both *miR*-*430* and *klf17* were strongly downregulated genes (*strongly down*). H3K27ac and H3K18ac were not affected in MZ*nps* embryos at *actb1*, whose transcription was unaffected (*unaffected*). (D) Line plots showing the average H3K27ac and H3K18ac signal at promoter and active enhancer regions in the different categories of zygotic genes presented in Figure 5D. (E) Boxplots comparing the change in H3K27ac and H3K18ac level at promoters (H3K27ac: strongly down vs mildly down: *P* = 1.1×10^−47^; mildly down vs unaffected: *P* = 7.0×10^−15^; strongly down vs unaffected: *P* = 1.7×10^−71^; H3K18ac: strongly down vs mildly down: *P* = 3.0×10^−46^; mildly down vs unaffected: *P* = 1.6×10^−18^; strongly down vs unaffected: *P* = 2.9×10^−70^, two-sample t-test) and active enhancers (H3K27ac: strongly down vs mildly down: *P* = 6.0×10^−19^; mildly down vs unaffected: *P* = 9.0×10^−20^; strongly down vs unaffected: *P* = 5.9×10^−86^; H3K18ac: strongly down vs mildly down: *P* = 4.2×10^−16^; mildly down vs unaffected: *P* = 1.5×10^−18^; strongly down vs unaffected: *P* = 3.2×10^−81^, two-sample t-test) of zygotic genes in different categories. This suggests that reductions in H3K27ac and H3K18ac correlated with the loss of transcription in MZ*nps* embryos.

**Figure S6.**
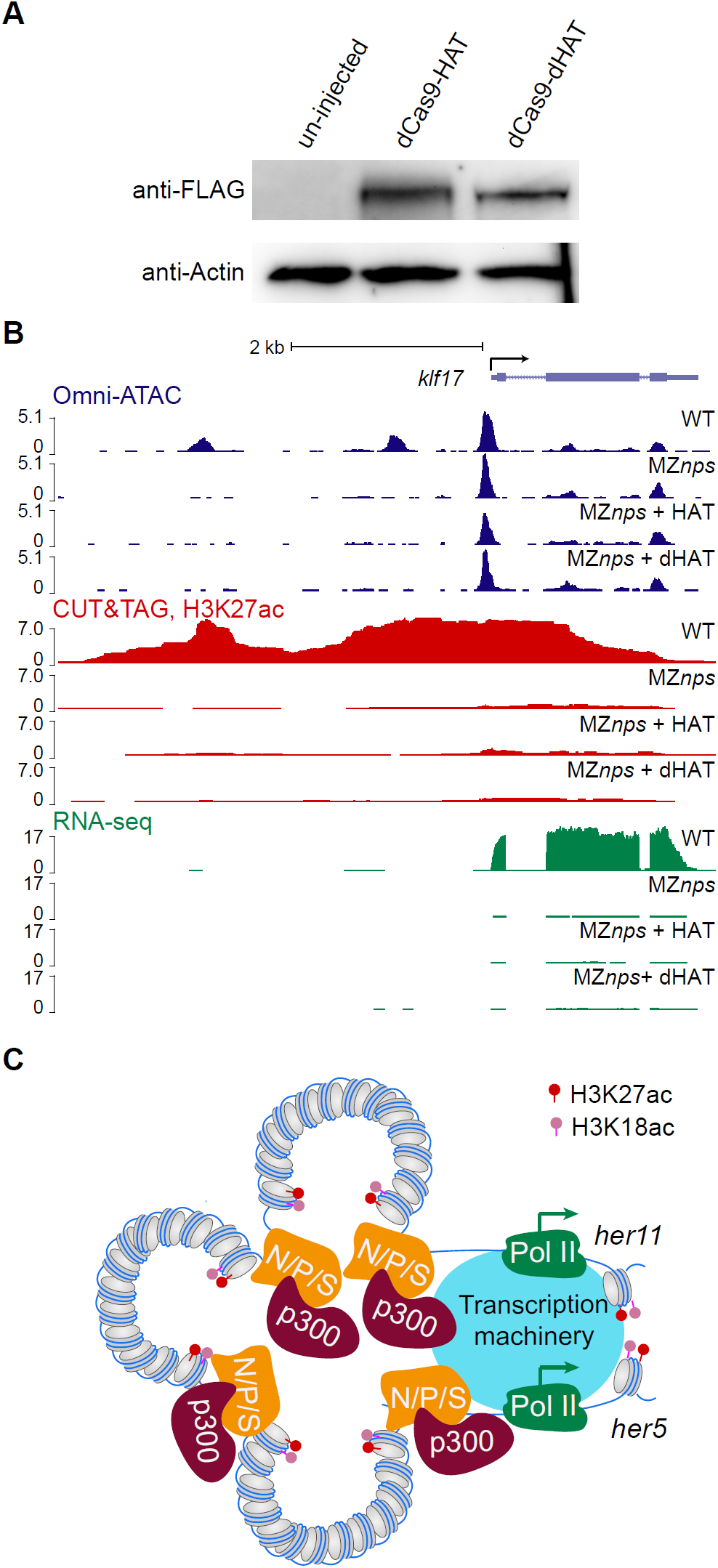
Restoration of H3K27ac partially rescue transcription of Nanog, Pou5f3, and Sox19b strong targets. (A) Immunoblotting was used to detect expression of dCas9-HAT (3xFLAG-dCas9-p300 Core (WT)) and dCas9-dHAT (3xFLAG-dCas9-p300 Core (D1399Y) in zebrafish embryos. (B) Genomic tracks of a control gene, *klf17*, that was not targeted by the dCas9-HAT system. H3K27ac and transcription levels were largely downregulated in MZ*nps* embryos and were not restored by dCas9-HAT in the absence of *klf17*-specific sgRNAs. Signal intensity in RPM (reads per million). (C) A model illustrating how NPS regulate H3K27ac and recruit Pol II to activate *her5* and *her11*, indicating that these two genes may share regulatory elements.

## Methods

### Zebrafish maintenance and generation of mutants

Fish lines were maintained in accordance with the AAALAC research guidelines following a protocol approved by the Yale University IACUC. Zebrafish husbandry and manipulation were performed as described (Howe et al., 2016; Westerfield, 2000). *nanog* and *sox19b* mutants were generated using CRISPR/Cas9 (Deltcheva et al., 2011; Haurwitz et al., 2010; Moreno-Mateos et al., 2015) targeting the following sequences: “AGCCCGGGTCTTGCGGGGCAGGG” and “GCAAACGGCTCGGCGCGGAGTGG,” respectively. A 4-nucleotide deletion mutant for *nanog* and a 5-nuleotide deletion mutant for *sox19b* were identified and maintained. See Table S1 for all oligos used to generate the mutants. MZ*nanog* embryos are not viable, as previously described (Gagnon et al., 2018; Veil et al., 2018). The *nanog* homozygous mutant line was maintained by injecting MZ*nanog* embryos with 25 pg of *nanog* mRNA (Lee et al., 2013) at the 1-cell stage. Consistent with published work (Pálfy et al., 2020), the *sox19b* homozygous mutants are viable. The *nanog* homozygous mutant was crossed with the *sox19b* homozygous mutant to obtain *nanog*^+/-^;*sox19b*^+/-^double heterozygotes, which were then crossed with *pou5f3*_hi349Tg/hi349Tg_ mutants (Burgess et al., 2002) to obtain *nanog*^+/-^;*pou5f3*^+/-^;*sox19b*^+/-^ triple heterozygotes. The *nanog*^+/-^;*pou5f3*^+/-^;*sox19b*^+/-^ triple heterozygotes were intercrossed and the resulting embryos were injected with 30 pg of *pou5f3* mRNA at the 1-cell stage. These embryos were raised and genotyped to obtain the *nanog*^-/-^;*pou5f3*^-/-^;*sox19b*^-/-^ triple homozygous mutant. To maintain the MZ*nps* mutant line, the *nanog*^-/-^;*pou5f3*^-/-^;*sox19b*^-/-^ triple mutants were intercrossed and the embryos were injected with 25 pg of *nanog* mRNA and 30 pg of *pou5f3* mRNA at the 1-cell stage. *sox19b* mRNA was not injected because the *sox19b* homozygous mutants are viable (Pálfy et al., 2020).

### sgRNA synthesis and injection

sgRNAs were designed and synthesized as previously described (Vejnar et al., 2016). Briefly, sgRNAs were designed using CRISPRscan (Moreno-Mateos et al., 2015). sgRNA templates were amplified using the oligos listed in Table S1 together with the following universal reverse primer: 5′ AAAAGCACCGACTCGGTGCCACTTTTTCAAGTTGATAACGGACTAGCCTTATTTTAACTTGCTATTTCTAGCTCT AAAAC-3′ (Vejnar et al., 2016). sgRNAs were then synthesized using the AmpliScribe T7-Flash Transcription Kit (Lucigen) and purified by ethanol precipitation or with RNeasy MinElute Cleanup Kit (QIAGEN). To generate mutants, 100 pg of Cas9 mRNA and 20 pg of sgRNA were injected into 1-cell stage embryos.

### Histone acetylation restoration in MZnps embryos with the dCas9-HAT system

The histone acetyltransferase domain from human p300 was subcloned from a published construct (Addgene, 61357) (Hilton et al., 2015) into the pT3TS-dCas9 plasmid to obtain pT3TS-dCas9-HAT. The HAT domain was inactivated by introducing a point mutation (D1399Y) in pT3TS-dCas9-HAT with the primers listed in Table S1 to produce pT3TS-dCas9-dHAT. dCas9-HAT mRNA and dCas9-dHAT mRNA were synthesized using the mMESSAGE mMACHINE T3 Transcription Kit (Thermo Fisher Scientific) according to manufacturer’s instructions. For qPCR and RNA *in situ* hybridization following histone acetylation by dCas9-HAT, *asb11* and *her5* were targeted separately. Specifically, 200 pg of dCas9-HAT mRNA or dCas9-dHAT mRNA and 30 pg of each sgRNA targeting either *asb11* or *her5* (sgRNAs 1-3 for *asb11* or sgRNAs 1-4 for *her5*) were injected into 1-cell stage embryos. When measuring transcript level by qPCR, *asb11* and *her5* sgRNAs were used as control sgRNAs for each other. For high-throughput sequencing experiments, *asb11* and *her5* were targeted simultaneously. 200 pg of dCas9-HAT mRNA or dCas9-dHAT mRNA together with 15 pg of each sgRNA (sgRNAs 1-3 for *asb11* and sgRNAs 1-4 for *her5*) were injected into 1-cell stage embryos. Embryos were collected at 4 hpf for subsequent assays.

### RNA in situ hybridization

Template DNAs for antisense RNA probes were amplified from 6 hpf cDNA using primers containing T3-promoter sequence in the forward primer as listed in Table S1. Digoxigenin (DIG) labeled RNA probes were synthesized using T3 RNA Polymerase (Roche) and purified by ethanol precipitation. RNA *in situ* hybridization was performed as described (Giraldez et al., 2005). Briefly, MZ*nps* embryos were injected at the 1-cell stage for targeted histone acetylation. Uninjected WT, uninjected MZ*nps* embryos, and injected MZ*nps* embryos were fixed at 4 hpf with 4% paraformaldehyde (PFA) overnight at 4°C. Fixed embryos were washed three times with phosphate-buffered saline (PBS) and dehydrated with a methanol series (25%, 50%, 75%, and 100% methanol). Dehydrated embryos were stored at −20°C for at least 24 hours and then rehydrated with a reverse methanol series. Pre-hybridization and hybridization were performed at 65°C for 3 hours and overnight, respectively. Embryos were washed extensively and blocked for 3 hours at room temperature before anti-DIG antibody incubation overnight at 4°C. After antibody incubation, embryos were stained with BCIP/NBT. Staining was stopped by incubating the embryos with 4% PFA overnight at 4°C. Embryos were washed briefly, mounted with a glycerol series (30%, 50%, 70%, and 86%), and imaged in 86% glycerol with a Zeiss stereo Discovery.V12 microscope.

### RNA-seq and real-time quantitative PCR (qPCR)

Total RNA was extracted from embryos at 4 hpf using TRIzol reagent (Thermo Fisher Scientific) following the manufacturer’s instructions and was treated with DNase I to remove any DNA contamination. Poly(A)-selected RNA-seq libraries were prepared and sequenced with an Illumina HiSeq system at the Yale Center for Genome Analysis. For qPCR, the extracted RNA was reverse-transcribed into cDNA using SuperScript III Reverse Transcriptase (Thermo Fisher Scientific). qPCR was performed using Power Sybr Green PCR Master Mix (Thermo Fisher Scientific). The efficiency of all primer-sets used, listed in Table S1, was determined to be between 90%-110%. *asb11* and *her5* levels were normalized to *actb1* (*beta*-*actin1*) (Yuan et al., 2010). Data were collected from three biological replicates. The two-tailed unpaired t-test was used to evaluate whether the transcription level was significantly different between two groups.

### Nascent transcriptome capture with Click-iT-seq

Click-iT-seq was performed as previously described (Chan et al., 2019). Briefly, 50 pmol of EU (5-ethynyl uridine), from the Click-iT Nascent RNA Capture Kit (C10365), was injected into 1-cell stage embryos. Total RNA was extracted from 20 embryos at 4 hpf using TRIzol reagent (Thermo Fisher Scientific). Nascent RNAs were captured and cDNA was synthesized with the SuperScript VILO cDNA synthesis kit (11754050) following the manufacturer’s instructions (C10365). Libraries were prepared from cDNA following the dUTP protocol and Illumina TruSeq proto col, and were sequenced with an Illumina HiSeq system at the Yale Center for Genome Analysis.

### Immunofluorescence imaging and Click-iT RNA imaging

As previously described (Chan et al., 2019), 25 pg of dCas9-3xGFP mRNA together with two sgRNAs targeting *miR-430* (100 pg each, see Table S1 for the amplification primers) were injected into 1-cell stage embryos. Embryos were fixed with 4% PFA at 2.5 hpf (the 256-cell stage) to perform immunofluorescence against GFP to label the *miR-430* locus. To ensure that the loss of transcription at the *miR*-*430* locus in the MZ*nps* embryos can be detected by imaging, Click-iT RNA imaging was performed together with *miR*-*430* locus labeling. Specifically, WT and MZ*nps* embryos were injected with 25 pg of dCas9-3xGFP mRNA, 100 pg each of the two *miR*-*430* targeting sgRNAs, and 50 pmol of EU at the 1-cell stage and fixed with 4% PFA at 2.5 hpf. Fixed embryos were washed three times with 1X PBST (phosphate-buffered saline + 0.5% Triton X-100), dehydrated with a methanol series (25%, 50%, 75%, and 100% methanol), stored at −20°C for at least 2 hours, and then rehydrated with a reverse methanol series. Rehydrated embryos were blocked with 10% BSA (in PBST) for 2-3 hours and incubated with anti-GFP antibody (1:1000; ThermoFisher Scientific, A-11120) overnight at 4°C. After three washes with 1X PBST, nascent transcripts were labeled using the Click-iT RNA Alexa Fluor 594 Imaging Kit (C10330) following the manufacturer’s instructions. Then, the embryos were incubated with Alexa Fluor Plus 488 anti-Mouse secondary antibody (1:1,000; ThermoFisher Scientific, A32723). Embryos were stained with DAPI, mounted in ProLong Diamond Antifade Mountant (ThermoFisher Scientific, P36965), and imaged using confocal fluorescence microscopy (ZEISS LSM 880 or Leica TCS SP8). Images were processed using Image-J software (Schneider et al., 2012) and are displayed as maximumintensity Z-projections unless otherwise noted. To screen for histone modification(s), immunofluorescence was performed together with *miR*-*430* locus labeling. Briefly, embryos were injected and treated as described. For the primary antibody incubation, anti-GFP antibody (mouse) in combination with a second antibody (rabbit)—against a specific histone modification—was used. All antibodies were diluted 1:1000. Antibodies against H3K27ac (ab4729), H3K18ac (ab1191), and H4K8ac (ab15823) were used.

### Omni-ATAC

To reduce contamination by mitochondrial DNA, Omni-ATAC (Corces et al., 2017) was adapted to zebrafish based on published methods (Buenrostro et al., 2013; Corces et al., 2017; Liu et al., 2018). Briefly, at 4 hpf, five embryos were manually deyolked in PBS and resuspended in cold lysis buffer (10 mM Tris-HCl pH 7.4, 10 mM NaCl, 3 mM MgCl_2_, 0.1% NP40, 0.1% Tween-20, and 0.01% digitonin). The lysate was incubated on ice for 5 minutes, then 1 ml of dilution buffer (10 mM Tris-HCl pH 7.4, 10 mM NaCl, 3 mM MgCl2, 0.1% Tween-20) was mixed with the lysate by inverting several times and samples were centrifuged at 500 g for 10 minutes at 4. The supernatant was removed and the purified nuclei were resuspended in the transposition reaction mixture (25 μl 2 × TD Buffer, 2.5 μl Tn5 transposase, 22.5 μl Nuclease-Free water) and incubated for 30 minutes at 37°C. DNA was then purified with the Qiagen MinElute Kit (Qiagen, 28004). Libraries were prepared using NEBNext High-Fidelity 2X PCR Master Mix (NEB, M0541) with the following conditions: 72°C, 5 minutes; 98°C, 30 seconds; 15 cycles of 98°C, 10 seconds; 63°C, 30 seconds; and 72°C, 1 minute. Libraries were purified with Agencourt AMPureXP beads (Beckman Coulter Genomics, A63881) and sequenced with the Illumina NovaSeq 6000 System at the Yale Center for Genome Analysis.

### ChIP-seq

Chromatin immunoprecipitation (ChIP) was performed as described (Bogdanovic et al., 2013; Leichsenring et al., 2013; Nelson et al., 2014; Zhang et al., 2014). Briefly, embryos were dechorionated at the 1-cell stage. For each assay, 1000 embryos were fixed at 4 hpf with 1.9% PFA for 15 minutes, quenched with 0.125M glycine, washed with cold PBS, and then frozen with liquid nitrogen and stored at −80°C until use. Nuclei were purified and lysed in 100 µl nuclei lysis buffer (50 mM Tris–HCl pH 7.5, 10 mM EDTA, 1% SDS) and diluted with 200 µl IP dilution buffer (16.7 mM Tris–HCl pH 7.5, 167 mM NaCl, 1.2 mM EDTA, 0.01% SDS). Lysates were sonicated with a Bioruptor Pico sonication device (Diagenode) using two rounds of 15 cycles of 30 seconds ON and 30 seconds OFF, with 15 minutes on ice in between rounds. Then, 27.6 µl of 10% Triton X-100 was added to the sonicated chromatin before centrifugation at 14,000 rpm for 10 minutes at 4°C and 5% of the supernatant was taken as input. Protein G Dynabeads (Invitrogen, 10003D) were pre-bound to antibodies overnight at 4°C and washed three times with 0.5% BSA in 1X PBS. 25 µl of antibody-bound beads was added to the remaining supernatant and incubated overnight at 4°C. Samples were washed five times with RIPA wash buffer (50 mM HEPES pH 7.6, 1 mM EDTA, 0.7% DOC, 1% Igepal, 0.5 M LiCl) and two times with 1X TBS (Tris-buffered saline; 50 mM Tris pH 7.5, 150 mM NaCl). DNA–protein complexes were eluted with 660 µl elution buffer (50 mM NaHCO3, 1% SDS) at 65°C for 15 minutes with occasional vortexing. Input samples were thawed, and three volumes of elution buffer was added. NaCl was added to samples to a final concentration of 0.2 M and then the samples were reverse-crosslinked overnight at 65°C. Samples were treated with RNase A (final concentration: 0.33 µg/µl) for 2 hours and Proteinase K (final concentration: 0.2 µg/µl) for 2 hours before purification. Libraries were prepared following the Illumina TruSeq protocol and sequenced with the Illumina NovaSeq 6000 System at the Yale Center for Genome Analysis. Antibodies against H3K27ac, Eomesa, and RNA Polymerase II were ChIP-validated in zebrafish (Bogdanovic et al., 2013; Nelson et al., 2014; Zhang et al., 2014). For H3K27ac ChIP-seq, 4 µg of anti-H3K27ac (ab4729) antibody was used for each sample. For Eomesa ChIP-seq, 40 µl of anti-Eomesa antibody (a gift from Dr. Fiona Wardle) was used for each sample. For ChIP-seq against RNA Polymerase II, 4 µg of anti-RNA polymerase II CTD repeat YSPTSPS antibody [8WG16] from Abcam (ab817) and 4 µg from BioLegend (664912) were used for each sample. Because there are no antibodies available to perform ChIP-seq against zebrafish Pou5f3 or Sox19b, we overexpressed Myc-tagged Pou5f3 and Myc-tagged Sox19b and performed ChIP-seq using 4 µg of anti-Myc-tag antibody (ab9132) for each sample. In detail, a 6x Myc tag was cloned into to the N-terminus of Pou5f3 and C-terminus of Sox19b in the pCS2 backbone. The plasmids were linearized with NotI (NEB, R0189) and mRNA (*myc*-*pou5f3* and *sox19b*-*myc*) was synthesized with the mMESSAGE mMACHINE SP6 Transcription Kit according to the manufacturer’s instructions. WT embryos were injected with 20 pg of *sox19b*-*myc* mRNA or 30 pg of *myc*-*pou5f3* mRNA at the 1-cell stage. To ensure that the enrichment of precipitated DNA was not due to non-specific binding of the anti-Myc antibody, ChIP-seq using the anti-Myc antibody was also performed on uninjected WT embryos.

### CUT&TAG against H3K27ac

CUT&TAG was adapted from the published protocol (Kaya-Okur et al., 2019). Briefly, at 4 hpf, five embryos were manually deyolked in PBS and then dissociated by pipetting up and down in 400 µl Dig-wash Buffer (20 mM HEPES, pH 7.5; 150 mM NaCl; 0.5 mM Spermidine; 1× Protease inhibitor cocktail; 0.015% Digitonin). Cells were spun down (600 × g for 3 minutes at 4°C), washed briefly with Dig-wash Buffer, and then resuspended in 50 µl of H3K27ac antibody buffer (2 mM EDTA and a 1:50 dilution of the H3K27ac antibody in Dig-wash Buffer). Cells were incubated with primary antibody at room temperature for 2 hours with gentle rotation. Cells were spun down and washed with Dig-wash Buffer (3 × 5 minutes) to remove unbound antibody. Then, cells were resuspended in 50 µl of Dig-med Buffer (20 mM HEPES, pH 7.5; 300 mM NaCl; 0.5 mM Spermidine; 1× Protease inhibitor cocktail; 0.015% Digitonin) containing a 1:200 dilution of the pA-Tn5 adapter complex (∼0.04 µM) and incubated at room temperature for 1 hour with gentle rotation. After incubation, cells were spun down and washed with Dig-med Buffer (4 × 5 minutes) to remove unbound pA-Tn5 complexes. Then, cells were resuspended in 100 µl of Tagmentation buffer (10 mM MgCl2 in Dig-med Buffer) and incubated at 37°C for 1 hour. Tagmentation was stopped by adding 2.25 µl of 0.5 M EDTA, 5.5 µl of 10% SDS, and 1 µl of 20 mg/mL Proteinase K to the reaction. Next, DNA was extracted with 244 µl of AMPureXP beads (Beckman Coulter Genomics, A63881). Libraries were prepared with the NEBNext High-Fidelity 2X PCR Master Mix as in Omni-ATAC, purified with Agencourt AMPureXP beads, and sequenced with the Illumina NovaSeq 6000 System at the Yale Center for Genome Analysis.

### Omni-ATAC data processing and analysis

LabxDB seq (Vejnar and Giraldez, 2020) was used to manage our high-throughput sequencing data and configure our analysis pipeline. Raw paired-end Omni-ATAC reads were adapter trimmed using Skewer (Jiang et al., 2014) and mapped to the zebrafish GRCz11 genome sequence (Yates et al., 2019) using Bowtie2 (v2.3.4.1) (Langmead and Salzberg, 2012) with parameters ‘-X 2000, –no-unal’. Unpaired and discordant reads were discarded. The alignments were deduplicated using samtools markdup (Li et al., 2009). Reads mapped to the + strand were offset by +4 bp and reads mapped to the – strand were offset by −5 bp (Buenrostro et al., 2013). Only fragments with insert size <=100 nt (effective fragments) were used to determine accessible regions. Genomic tracks were created using BEDTools (Quinlan and Hall, 2010) and utilities from the UCSC genome browser (Lee et al., 2020). Fragment coverage on each nucleotide was normalized to the total number of effective fragments in each sample per million fragments. For genome-wide analysis, reads mapped to mitochondrial DNA and contigs were filtered out and only uniquely mapped reads (with alignment quality ≥ 30) were used.

### ChIP-seq data processing and analysis

Raw ChIP-seq reads were adapter trimmed, mapped, deduplicated, and tracks were created using the same method described in the previous section but using the default parameters for Bowtie2 for read mapping. For paired-end samples, fragment coverage on each nucleotide was normalized to the total fragments in each sample per million fragments. For single-read samples, reads were extended to the predicted fragment size from MACS2 (see Peak calling below) (Zhang et al., 2008) and read coverage on each nucleotide was calculated based on the extended reads and was normalized to the total number of reads in each sample per million reads. For genome-wide analysis, reads mapped to mitochondrial DNA and contigs were filtered out and only uniquely mapped reads (with alignment quality ≥ 30) were used.

### Peak calling

Peaks were called using MACS2 (Zhang et al., 2008) for Omni-ATAC and ChIP-seq data. For Omni-ATAC, peaks were called with the additional parameters ‘-f BEDPE –nomodel –keep-dup all’ with significance cutoff at *P* = 10^−8^. To determine high-confidence accessible regions, the three WT Omni-ATAC replicates were merged, and then peaks were called from the merged data. Only those peaks called from merged data that overlap with peaks called from all three biological replicates individually were defined as accessible regions. For ChIP-seq of transcription factors, narrow peaks were called using MACS2 with the additional parameters ‘-f BEDPE –nomodel –keep-dup all’ for paired-end samples and ‘-f BAM –keep-dup all’ for single-read samples. Peaks from ChIPseq of Nanog, Pou5f3, and Sox19b were called with the default significance cut-off (q = 0.05). To identify NPS-dependent and NPS-independent regions, peaks called on Eomesa ChIP-seq in MZ*nps* embryos were compared to peaks called by merging Eomesa ChIPseq data from MZ*nps* and WT embryos. Peaks from merged data that overlap with peaks in MZ*nps* were classified as NPS-independent regions, all remaining peaks from merged data were classified as NPS-dependent regions. Peak calling was performed using a significance level of *P* = 10^−5^ on the merged set while *P* = 10^−3^ was used otherwise. For ChIP-seq of histone modifications, broad peaks were called with the additional parameters ‘-f BEDPE –nomodel –keep-dup all-q 0.05 –broad –broad-cutoff 0.05’ to identify peaks for broad regions.

### Differential accessibility analysis

To identify regions where accessibility was significantly changed in MZ*nps* embryos, the fragment coverage of each accessible region of the three WT replicates was compared with that of the three MZ*np*s replicates using DESeq2 (Love et al., 2014) with false discovery rate (FDR) < 0.01. Accessible regions with significantly lower Omni-ATAC signal in MZ*nps* embryos were further sub-divided into two categories: regions with no overlap with peaks called at a low significance cutoff (*P* = 10^−3^) from any MZ*nps* Omni-ATAC replicate were classified as Group 1 (accessibility completely lost); and the other regions were classified as Group 2 (accessibility decreased but still accessible). Additionally, accessible regions where accessibility was not significantly changed and the fragment coverage difference between embryos WT and MZ*nps* embryos was less than 30% were classified as Group 3 (accessibility unaffected).

### Definition of promoters and enhancers

Promoters were defined as the region within 500 bp of a transcription start site (TSS) of annotated transcripts from Ensembl v92 (Yates et al., 2019) and Cap Analysis of Gene Expression (CAGE) datasets (Nepal et al., 2013) at 3.3 hpf, 3.7 hpf, 4 hpf, and 6 hpf. The promoter of the most abundant transcript isoform, as determined by the CAGE data, was used to define the promoter of a gene. If no isoform had a maximum CAGE signal ≥ 0.5 RPM, the promoter was defined using a gene model that was constructed by merging all transcript isoforms annotated from Ensembl. Enhancers were defined as accessible regions outside of the promoter regions (TSS ± 500 bp) and overlapping with H3K27ac region (H3K27ac peak region with 200 nt extension on both sides). Enhancers within 4,500 bp of the promoter of a zygotic gene (see below) were considered to be associated with the gene. If an enhancer was within 4,500 bp of the promoters of multiple genes, the enhancer was considered to be associated with the gene closest to it. Accessibility of each enhancer was defined as the sum of Omni-ATAC signal across all nucleotides within the enhancer region. The H3K27ac signal of each enhancer was defined as the sum of H3K27ac signal across all nucleotides within the flanking 500 bp region (the region that is outside of the enhancer but with a distance ≤ 500 bp to the enhancer region). The H3K27ac signal of promoters was defined as the sum of H3K27ac signal across all nucleotides within the flanking 500 bp region of the H3K27ac-overlapping accessible region that also overlapped with the promoter region.

### RNA-seq processing and analysis

Raw reads were aligned to the zebrafish GRCz11 genome sequence using STAR 2.7.5b (Dobin et al., 2012) with the parameters ‘–alignEndsType Local –sjdbScore 2’. Genomic sequence index for STAR was built including exon-junction coordinates from Ensembl v92. Gene models were constructed by merging the transcript isoforms of each gene. Reads overlapping at least 10 nucleotides of the gene annotation were considered to be mapped to the gene. For read counts of the genes, each locus where a read was mapped was assigned a weight equal to 1 divided by the total number of loci to which the read was mapped. Read counts per gene were sum of the weights assigned to the gene. For miR-430, reads overlapping the region from coordinate 28,693,371 to 28,709,534 on chromosome 4 were counted as miR-430 cluster reads. Read counts from regular RNA-seq experiments were normalized to the total number of reads mapped to the zebrafish genes per million. For Click-iT-seq experiments, read counts were normalized by the total number of mitochondrial reads mapped to mitochondrial protein coding genes. Zygotic genes were determined using a previously described method (Chan et al., 2019). Briefly, a zygotic gene was defined as a gene identified by (Heyn et al., 2014; Lee et al., 2013) as a zygotic gene, or a gene with a nascent transcription level ≥ 10 RPKM in exon regions and at least 4-fold more nascent transcription in WT than α-amanitin-treated embryos in exon or intron regions.

### Motif enrichment analysis

Motifs for Nanog, Pou5f3, Sox19b in zebrafish were identified using DREME in MEME suite (Bailey, 2011) based on all peaks called from the zebrafish ChIP-seq datasets at a significance cutoff at *P* = 10^−5^. Due to the high co-occupancy of Nanog and Pou5f3 and the similarity of their binding motifs, a second Nanog binding motif (Nanog motif 2) was identified from regions bound by Nanog that do not overlap with the binding regions of Pou5f3 and Sox19b. Pou5f3 motif, Sox19b motif, and both Nanog motifs were used to determine the contribution of the sequence context to the pioneering activities of different transcription factors. Motif enrichment was performed using AME in MEME suite (McLeay and Bailey, 2010) with default parameters on all known transcription factor binding motifs from the Motif database on the MEME suite website (http://memesuite.org/doc/download.html) and HOMER website (http://homer.ucsd.edu/homer/custom.motifs). Motifs for human and mouse transcription factors were used as the motifs for their homologous transcription factors in zebrafish. Homologs between zebrafish and human and mouse were identified using BioMart on Ensembl genome browser (Yates et al., 2019). Only transcription factors (or transcription factor homologs in zebrafish) with mRNA translation rate ≥ 5 RPKM at 2 hpf in zebrafish embryos (Chan et al., 2019) were included in this analysis. The same sample size (either 1000 or the minimum sample size among the groups) was used for motif enrichment across different conditions to generate comparable significance levels between conditions. Transcription factors were classified as housekeeping genes if they or their homologs were reported as housekeeping genes (Eisenberg and Levanon, 2013). Transcription factors (or their homologs) not listed in (Eisenberg and Levanon, 2013) were classified as housekeeping genes or developmental genes according to (Murphy et al., 2018).

### Motif frequency, motif strength, and nucleosome occupancy

The motif frequency within each region was determined by the number of sequences matching the binding motif of the specific transcription factors using FIMO (Grant et al., 2011). Motif co-occurrence was determined if a region contains sequences that match the binding motifs of both transcription factors. Motif strength, also referred to as motif score in this paper, was calculated as -log10 of the corresponding significance (*P* value) of the sequence that matches the transcription factor binding motif from FIMO. If a region contained multiple sequences matching the transcription factor binding motif, the highest motif score was used for that region. Nucleosome occupancy scores at nucleotide resolution were predicted using the model from (Kaplan et al., 2009).

### Heatmaps, plots, and networks

Heatmaps based on Omni-ATAC and ChIP-seq data were created using R 3.6 and the pheatmap package (https://github.com/raivokolde/pheatmap). Motif overrepresentation heatmaps were created using the R package gplots (https://github.com/talgalili/gplots). Box plots, bar plots, and violin plots were created using the R package ggplot2 (Wickham, 2016). The network of transcription factor co-occurrence was created using the R package igraph (Csardi and Nepusz, 2006). Biplots and line plots were created using Python 3.8 and the Matplotlib library (Hunter, 2007).

### Quantification and statistical analysis

For qPCR, statistical tests were performed on the data from three biological replicates using a two-tailed unpaired t-test. Fisher’s exact test was performed to calculate the significance of co-binding of different factors in Venn diagrams and the difference in percentage between different groups. Wilcoxon rank-sum test was used to calculate the significance of motif scores between different rescue conditions. The two-sample t-test was performed to calculate the significance between different groups for the rest of this paper. Pearson correlation was used to represent the correlation between two variables.

